# Transcript- and annotation-guided genome assembly of the European starling

**DOI:** 10.1101/2021.04.07.438753

**Authors:** Katarina C. Stuart, Richard J. Edwards, Yuanyuan Cheng, Wesley C. Warren, David W. Burt, William B. Sherwin, Natalie R. Hofmeister, Scott J. Werner, Gregory F. Ball, Melissa Bateson, Matthew C. Brandley, Katherine L. Buchanan, Phillip Cassey, David F. Clayton, Tim De Meyer, Simone L. Meddle, Lee A. Rollins

## Abstract

The European starling, *Sturnus vulgaris*, is an ecologically significant, globally invasive avian species that is also suffering from a major decline in its native range. Here, we present the genome assembly and long-read transcriptome of an Australian-sourced European starling (*S. vulgaris* vAU), and a second North American genome (*S. vulgaris* vNA), as complementary reference genomes for population genetic and evolutionary characterisation. *S. vulgaris* vAU combined 10x Genomics linked-reads, low-coverage Nanopore sequencing, and PacBio Iso-Seq full-length transcript scaffolding to generate a 1050 Mb assembly on 1,628 scaffolds (72.5 Mb scaffold N50). Species-specific transcript mapping and gene annotation revealed high structural and functional completeness (94.6% BUSCO completeness). Further scaffolding against the high-quality zebra finch (*Taeniopygia guttata*) genome assigned 98.6% of the assembly to 32 putative nuclear chromosome scaffolds. Rapid, recent advances in sequencing technologies and bioinformatics software have highlighted the need for evidence-based assessment of assembly decisions on a case-by-case basis. Using *S. vulgaris* vAU, we demonstrate how the multifunctional use of PacBio Iso-Seq transcript data and complementary homology-based annotation of sequential assembly steps (assessed using a new tool, SAAGA) can be used to assess, inform, and validate assembly workflow decisions. We also highlight some counter-intuitive behaviour in traditional BUSCO metrics, and present Buscomp, a complementary tool for assembly comparison designed to be robust to differences in assembly size and base-calling quality. Finally, we present a second starling assembly, *S. vulgaris* vNA, to facilitate comparative analysis and global genomic research on this ecologically important species.

## 1. Introduction

The European starling (*Sturnus vulgaris*) is a globally invasive passerine that was deliberately introduced during early European acclimatisation efforts into North America, Australia, New Zealand, and South Africa during the mid-late 19^th^ century (Feare 1985). More recently, the species was accidentally introduced into South America (Palacio et al. 2016). Since these introductions the invasive ranges of the starling have been expanding, with the species now occupying a range in excess of 38,400,000 km^2^ globally (BirdLife International 2020), posing threats to the economics and health of the agriculture industry, as well as local biodiversity (Bomford & Sinclair 2002; Koch et al., 2009; Palacio et al., 2016; Linz et al., 2017). Recent molecular ecology studies of individuals from the invasive ranges of North America, Australia, and South Africa report that these populations are undergoing rapid and independent evolution in response to novel local selection pressures (Phair et al., 2018; Hofmeister et al., 2019; Bodt et al., 2020; Stuart & Cardilini et al., 2020), a common phenomenon in many invasive populations (Prentis et al., 2008). This suggests the starling has a flexible invasion strategy, potentially enabling colonisation of ecosystems vastly different from those in their native range.

Despite their invasive range success, European starlings are increasingly of ecological concern within their native range (Rintala et al., 2003; Robinson et al., 2005). High densities of native range starlings have traditionally been supported by cattle farming across Europe, because starlings preferentially feed in open grasslands, and benefit from invertebrates in overturned soil produced by livestock grazing (Coleman 1977). A shift towards modern indoor cattle rearing processes across Europe may contribute to the decline in starling numbers, which has been a concern since the 1980s (Wretenberg et al., 2006). This decline is reflected globally, with starling and other avifauna numbers decreasing sharply over the last few decades (Spooner et al., 2018; Rosenberg et al., 2019), though this may be further amplified for starling populations subjected to control strategies to reduce their economic impact (Linz et al., 2007). The biological and ecological importance of this species is evident from its prolific use in research, as it is the most studied non-domesticated passerine (Bateson & Feenders 2010). It is evident that future research on the European starling will focus on identifying patterns of evolutionary diversification, and investigating genes associated with invasion success. Such research provides important information for the improvement of control measures and may also provide insight into recovery and dispersive potential in other species that would benefit global conservation efforts. For this, a high-quality, annotated reference genome is essential.

Once reliant on large consortia, assembling high-quality reference genomes for genetic analyses is now commonplace. Nevertheless, *de novo* assembly of non-model organism genomes still poses many challenges. Best practices may have not been established for the study species/data, and basic information such as genome size, repeat landscape, and ploidy may be unknown. Furthermore, high-quality references can be generated in multiple ways, which can serve varied research purposes. Rapid developments in both sequencing technology and bioinformatics methods can quickly outdate benchmarking attempts. Whilst not always documented in final publications, the standard practice for non-model species genomes is to select from multiple assemblies generated using different assembly methods, none of which is universally best (Rhie et al., 2020; Whibley et al., 2020). This complexity can be magnified further when sequencing occurs across multiple technology platforms that may be combined and utilised in different ways (Jayakumar & Sakakibara 2019; Kono & Arakawa 2019). The challenge is then to select the best combination of tools and assembly decisions, based on the quality of the genome assemblies produced.

A multitude of tools and approaches are available for genome assembly assessment, though some may not be applicable or feasibly implemented for a particular species/assembly and/or the data available (e.g. Bradnam et al., 2013; Hunt et al., 2013; Yuan et al., 2017). Common approaches employed to guide genome assembly decisions focus on contiguity (how continuous the assembled sequences are) and completeness (whether the assembly contains all the genetic information for that species). Two such approaches are assembly statistics (e.g., contig/scaffold counts, and L50/N50 statistics of the number and shortest length of sequences needed to cover 50% of the assembly) and “Benchmarking Universal Single Copy Orthologs” (BUSCO) estimates of genome completeness (Simão et al., 2015). Assembly statistics are very quick to generate and easy to understand, but interpretation can be challenging due to hidden assembly errors and artefacts, which can create false signals. BUSCO assesses the presence or absence of highly conserved lineage-specific genes but is limited to a set of common single-copy genes that may represent easier regions of the genome to sequence and assemble based on current bioinformatics technologies. Furthermore, BUSCO analysis is vulnerable to unpredictable misreporting of presence/completeness for specific genes as a consequence of assembly differences elsewhere in the genome (Edwards et al., 2018; Edwards 2019; Field et al., 2020; Edwards et al., 2021). In addition to the above drawbacks, these methods do not explicitly test the genome assembly’s ability to perform the role for which it was intended (e.g., to serve as a reference genome for specific genomic analysis).

Here, we present the first official release of the European starling draft genome, *S. vulgaris* vAU. This assembly represents the first synthesis of species-specific full-length transcripts, together with genomic data for this species. In this paper, we complement genome statistics and BUSCO completeness with transcriptome-and annotation-based assessments that help determine genome assembly quality and completeness in the absence of a reference genome to benchmark against. We demonstrate how full-length transcripts can be utilised in genome assembly scaffolding and assessment, in addition to transcriptome construction and annotation. We show how BUSCOMP (https://github.com/slimsuite/buscomp) can help avoid over-interpretation or misinterpretation of small differences in BUSCO completeness. We also explore how lightweight homology-based annotation by GeMoMa (Keilwagen et al., 2018), can be used as an assembly assessment using a new tool, SAAGA (https://github.com/slimsuite/saaga). Finally, we compare the Australian *S. vulgaris* vAU assembly (GCF_ JAGFZL000000000) with an additional (short read) assembly of a North American bird, *S. vulgaris* vNA (GCF_001447265.1), enabling reference-specific biases to be identified in future starling genomics studies.

## 2. Material and Methods

### 2.1 Genome assembly and scaffolding

The *S. vulgaris* vAU genome assembly used 10x Chromium linked reads and low coverage ONT long reads (Appendix 1: Genomic DNA sample collection, gDNA extraction, and sequencing), and was produced via eight assembly steps (Fig. 1). The 10x reads were assembled into an initial diploid assembly using supernova (v2.1.1) (Weisenfeld et al., 2017) with barcode fraction and reads subsample calculated following supernova best practices for a genome size based on k-mer counts calculation by Jellyfish v2.2.10 (Marçais & Kingsford 2011) (parameters: bcfrac = 0.8, maxreads = 550 million, Supplementary Materials: Appendix 2, Validation of Supernova genome size prediction using Jellyfish, Supplementary Materials: Fig. S1). This assembly was then split into non-redundant primary and alternative haploid assemblies using Diploidocus (parameters: runmode= diphapnr) (v0.9.5) (https://github.com/slimsuite/diploidocus). First, both Supernova pseudohap2 assemblies were combined and any sequences lacking definitive base calls (100% Ns) were removed. Remaining scaffolds were size-sorted and gaps reduced in size to a maximum of 10 Ns then subject to an all-by-all search with minimap2 (v2.17) (Li 2018) (--cs -p 0.0001 -x asm20 -N 250). (Note that gap size reduction is used for minimap2 searching only, and the non-redundant pseudodiploid assembly produced has the same gap sizes as generated by Supernova.). Any sequences that were 100% contained within another sequence were removed. Where two or more scaffolds had an 100% identical sequence, only one was kept. Scaffolds are then matched into haplotig pairs based on their Supernova names. Where a single haplotig is found, it is assigned as diploid, under the assumption that the two original haplotigs were identical with one removed, and added to the primary assembly. (Note: it is possible that only one parent had this scaffold, e.g., a sex chromosome scaffold or structural variant.). If two haplotigs are identified, the longest is assigned to the primary assembly and the shorter to the alternative assembly. The primary assembly should therefore contain an entire haploid copy of the genome, whilst the alternative assembly contains the subset of scaffolds with heterozygous haplotigs.

**Figure 1:**
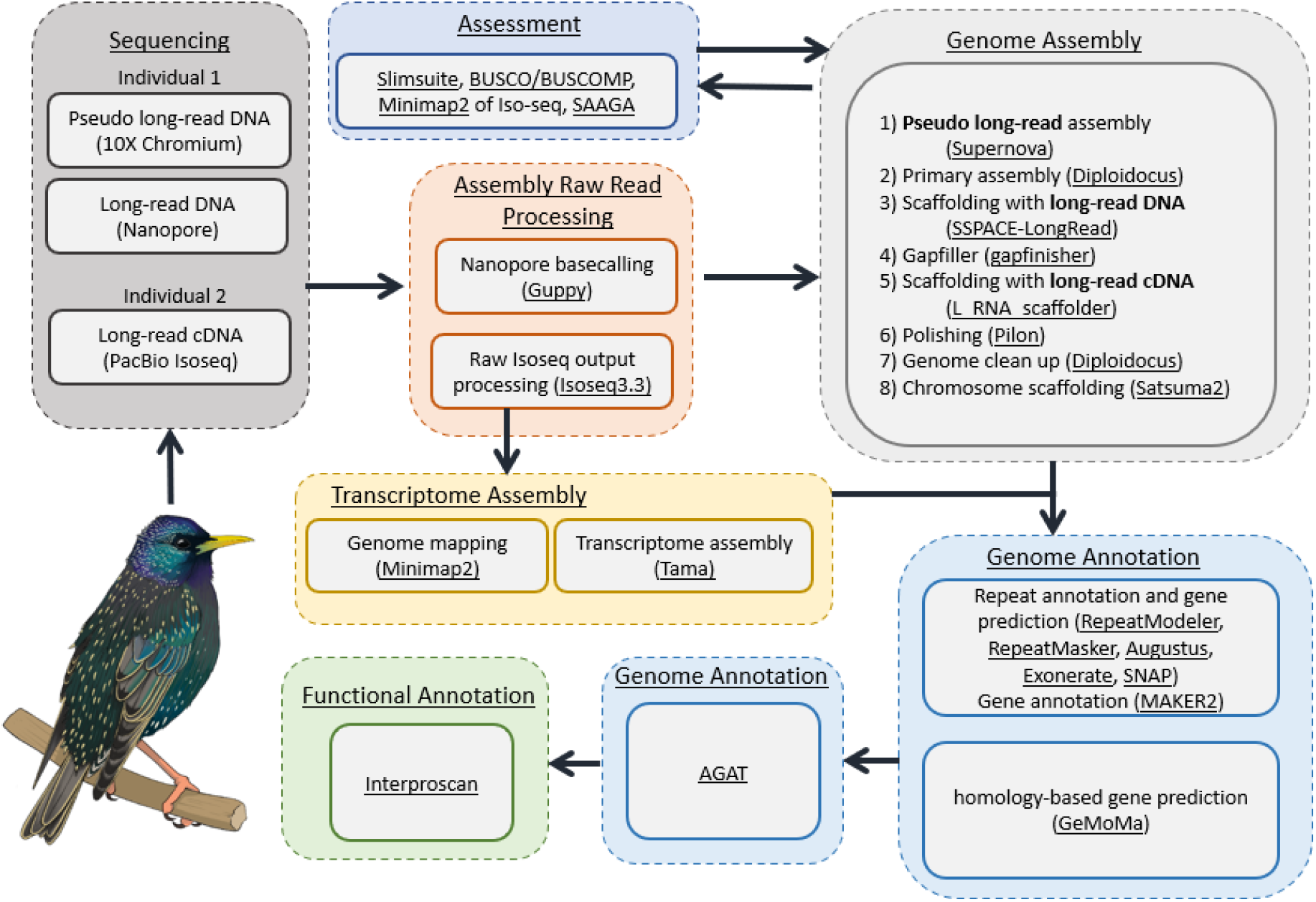
Workflow for genome assembly and annotation. A summary of all the experimental methods used for sequencing, genome assembly, transcriptome assembly, genome annotation, and functional annotation, with programs used underlined.

The primary haploid assembly produced by Diploidocus was scaffolded using the filtered ONT reads using the program SSPACE-LongRead (v1-1) (Boetzer & Pirovano 2014). The filtered ONT reads were then used to gap-fill the assembly using Gapfinisher (v1.0) (Kammonen et al., 2019). Clustered high-quality Iso-Seq reads (see section 2.2 cDNA analysis) were then used for a secondary round of scaffolding using L_RNA_scaffolder (Xue et al., 2013). Paired-end 10x linked reads were processed with 10X Genomics long ranger (v2.2) and mapped onto this scaffolded assembly using bwa *mem* before error correction of SNPs and indels using Pilon (v1.23) (Walker et al., 2014) (parameters: --diploid –fix all settings). To validate the scaffolds, the assembly was analysed using the Break10x toolkit in Scaff10x (v3.1) (https://github.com/wtsi-hpag/Scaff10X). The assembly was further checked for assembly artefacts and contamination using Diploidocus (parameters: runmode=purgehaplotig & runmode=vecscreen (ref); runmode=DipCycle was tested yet discarded due to over-pruning, see Supplementary Materials: Fig. S2) (v0.9.5). Avian species are characterised by distinctive and constrained karyotypes, generally comprised of approximately 10 macrochromosomes and approximately 30 indistinguishable microchromosomes (Griffin et al., 2007; O’Connor et al., 2019), a pattern to which the *S. vulgaris* genome conforms (Calafati & Capanna 1981). Therefore, we aligned our assembly to the chromosome scale assembly of zebra finch (Taeniopygia guttata) (NCBI= GCF_008822105.2) (Balakrishnan et al., 2010) using Satsuma2 (https://github.com/bioinfologics/satsuma2) to create putative chromosomes assuming orthology. This assembly formed the final updated draft genome we present for the species: *Sturnus vulgaris* vAU.

### 2.2 Transcriptome assembly and analysis

Raw PacBio Iso-Seq whole transcript reads (Appendix 3: Transcriptome sample collection, RNA extraction, and sequencing) were processed using the protocol outlined in SMRT Link (v9.0) (PacBio, California, United States). Briefly, this involved generating Circular Consensus Sequences (CCS) using CCS (v4.2.0), which were then processed using Lima (v1.11.0) for primer removal and demultiplexing. The reads were further processed (PolyA tail minimum length = 8) and clustered using Iso-Seq (v3.3). The high-quality clustered Iso-Seq reads were then aligned to the reference genome (see section 2.1 Genome assembly and scaffolding) using minimap2 (v2.17) (Li 2018), before further processing using Tama collapse (Kuo et al., 2020) (settings -a 100 -z 30 -sj sj_priority -lde 5). Both these steps were assessed using BUSCO (v3.0.2b) (Simão et al., 2015) (parameters: aves lineage, transcriptome mode), alongside a short read transcriptome produced from *S. vulgaris* liver RNA (Richardson et al., 2017), as well as other available avian Iso-Seq transcriptomes (Workman et al., 2018; Yin et al., 2019).

### 2.3 Genome annotation and functional annotation

Each stage of genome assembly was annotated using Gemoma v1.7.1 (Keilwagen et al., 2018) using the 26 avian genome annotations available on Ensembl at the time this analysis was conducted (Supplementary Materials, Table S1) and with the high-quality clustered Iso-seq, as RNA evidence. The gemoma *GeMoMaPipeline* function was run to complete the full pipeline with a maximum intron size of 200 kb (parameters: tblastn=false GeMoMa.m=200000 GeMoMa.Score=ReAlign AnnotationFinalizer.r=SIMPLE pc=true o=true).

The final *S. vulgaris* vAU genome assembly was also annotated with maker2 (Holt & Yandell 2011) (blast+ v2.9 (Camacho et al., 2009), AUGUSTUS v3.3.2 (Stanke & Morgenstern 2005), EXONERATE v2.2.0 (Gs & E 2005), REPEATMASKER v4.0.7 (Smit et al., 2013), REPEATMODELER v1.0.11 (Flynn et al., 2020), and SNAP v0.15.4 (Korf 2004) using repeat-filtered Swiss-Prot protein sequences (downloaded Aug 2018) (UniProt Consortium 2019). A custom AUGUSTUS species database was created by running BUSCO using the Optimization mode Augustus self-training mode (--long), using the aves database for lineage. MAKER2 was run using the recommended protocol, including generation of a repeat library, and with the TAMA-processed Iso-Seq data included as primary species transcript evidence, and the pre-existing short read liver transcript data (Richardson et al., 2017) provided as alternate transcript evidence in the first iteration of the MAKER2 annotation process. We ran MAKER2 for a total of three training runs, using the hidden Markov models (HMMs) produced from SNAP training in each subsequent run. *Ab initio* genes were not retained in the final annotation model to produce high-quality and conservative gene predictions. GeMoMa and Maker2 annotations for the final *S. vulgaris* vAU assembly were combined using the agat *agat_sp_merge_annotations* function to produce the final annotation. Functional annotation of protein-coding genes were generated using Interproscan 5.25–64.0 (parameters: -dp - goterms -iprlookup -appl TIGRFAM, SFLD, Phobius, SUPERFAMILY, PANTHER, Gene3D, Hamap, ProSiteProfiles, Coils, SMART, CDD, PRINTS, Pro SitePatterns, SignalP_EUK, Pfam, ProDom, MobiDBLite, PIRSF, TMHMM). blast was used to annotate predicted genes using all Swiss-Prot proteins (parameters: -evalue 0.000001 -seg yes - soft_masking true -lcase_masking -max_hsps). Annotation summaries were generated using the agat *agat_sp_functional_statistics*.*pl* script, bedtools was used to calculate gene coverage statistics. Gene ontology terms were assigned using wego v2.0 (Ye et al., 2018).

### 2.4 Annotation assessment using SAAGA: Summarise, Annotate & Assess Genome Annotations

SAAGA (Summarise, Annotate & Assess Genome Annotations) (v0.5.3) (https://github.com/slimsuite/saaga) was used to assess annotation quality and compare predicted proteins to the repeat- and transposase-filter Swiss-Prot protein sequences used for maker2 annotation (above). SAAGA performs a reciprocal MMseqs2 (Steinegger & Söding 2017) search of annotated proteins against a (high-quality) reference proteome, identifying best hits for protein identification and employing coverage ratios between query and hit proteins as a means of annotation assessment to generate summary statistics, including:

- **Protein length ratio**. The length ratio of the annotated proteins versus its top reference hit
- **F1 score**. An annotation consistency metric calculated using the formula:

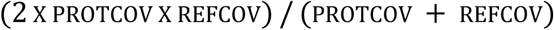

where protcov is the proportion of the annotated protein covered by its best reference protein hit, and refcov is the proportion of the best reference protein hit covered by the annotated protein.
- **Completeness**. The summed percentage coverage of reference proteome.
- **Purity**. The summed percentage reference coverage of the annotated proteome.
- **Homology**. The percentage of annotated genes with any hit in reference.
- **Orthology**. The percentage of annotated genes with reciprocal best hits in reference.
- **Duplicity**. The mean number of annotated genes sharing the same best reference hit.
- **Compression**. The number of unique annotated genes that were the top hit for reference proteins, divided by the total number of reference proteins with a hit.
- **Multiplicity**. The ratio of total number of annotated genes to reference proteins.

For protein length ratio and F1 score, values close to 1 means that the query protein closely matches the length of the hit protein, indicating high fidelity of the gene prediction model and underlying assembly. The remaining metrics will be closer to 1 (or 100%) for complete annotations and assemblies without duplications, akin to BUSCO scores. Although the maximum achievable value for these metrics will generally be unknown, comparative values can be used to assess improvement in assembly and/or annotation.

SAAGA scores may be used to compare alternate annotations of the same assembly, or to compare alternative assemblies in conjunction with consistent annotation. Low genome contiguity, misassembles, or frameshifting indels will affect the quality of predicted genes, with poorer assemblies reporting more fragmented or truncated genes. This approach has been facilitated by the rapid homology-based gene prediction program GeMoMa, which uses reference genome annotation to predict protein-coding genes in the target genome. The program can be run from one line of code and may be parallelised to run much faster than other annotation software (e.g., Maker2). The ease of this annotation tool opens the way for conducting annotations for the purpose of assessment on sequential or even competing genome annotation steps. Assessing the quality of protein-coding region predictions will help ensure the final genome assembly can produce a high-quality annotation. Here, we used the repeat-filtered Swiss-Prot database used in annotation, and the *Gallus gallus* reference proteome (UP000000539_9031), to assess predicted protein quality and annotated proteome completeness.

### 2.5 Genome assembly completeness assessment

Assembly contiguity and completeness was assessed for sequential genome assembly steps of the *S. vulgaris* vAU assembly and compared to existing passerine chromosome level assemblies available on NCBI, including the *S. vulgaris* vNA assembly (Assembly accession GCF_001447265.1, Supplementary Material: Appendix 4, Assembly and annotation of the *S. vulgaris* vNA genome version).

#### 2.5.1 BUSCO and BUSCOMP assembly completeness assessment

Genome completeness was estimated using BUSCO (v3.0.2b, genome mode, aves lineage). BUSCO resulted were collated across all assemblies using BUSCOMP v0.10.1 (https://github.com/slimsuite/buscomp). BUSCOMP collated BUSCO outputs across all genome assembly stages and compiled a maximal non-redundant set of 4727 complete BUSCOs found at single copy in at least one assembly. Compiled BUSCO predicted gene sequences were mapped onto each assembly to be rated with minimap2 v2.17 (Li 2018) and re-scored in terms of completeness, thereby providing a robust and consistent means of assessing comparable completeness across assemblies of the same genome.

#### 2.5.2 PacBio Iso-Seq completeness assessment

The PacBio Iso-Seq reads were mapped on to genome assemblies using minimap2 (parameters: -ax splice -uf --secondary=no --splice-flank=no -C5 -O6,24 -B4) (Li 2018) and the number of Iso-Seq transcripts mapping on to each assembly, and their corresponding mapping quality, was calculated.

#### 2.5.3 KAT k-mer completeness assessment

The final genome assembly completeness was assessed by examining the read k-mer frequency distribution with different assembly copy numbers based on the 10x Chromium linked reads using K-mer Analysis Toolkit (kat) v2.4.2 (Mapleson et al., 2017) (30 bp trimmed for R1 reads, and 16 bp trimmed for R2 reads).

### 2.6 Additional genome statistics

The Iso-Seq and final annotation transcript density, final annotation gene density, global SNP variant density (based on a whole genome data set of 24 individuals from United Kingdom, North America, and Australia, N=8 (Hofmeister *et al*. 2021), and GC-content were calculated in sliding windows of width 1 Mb using bedtools v 2.27.1 (Quinlan & Hall 2010), and plotted across the largest 32 scaffolds in our final genome assembly (representing more than 98% of the total assembly captured on putative chromosomes orthologous to other avian chromosomes) using circlize (v 0.4.9) (Gu et al., 2014).

### 2.7 Genome assembly correction

NCBI VecScreen flagged possible bacterial and adapter contamination in the final *S. vulgaris vAU* assembly, which was missed by earlier contamination screening steps. An updated version of Diploidocus (runmode vecscreen) was run to mask shorter adapter sequences and flag additional organism contaminates (screenmode=purge vecmask=27). Four related bacterial strains (Delftia acidovorans SPH-1, Acidovorax sp. JS42, Alicycliphilus denitrificans K601, Paraburkholderia xenovorans LB400) were identified, and so GABLAM v2.30.5 (Davey et al., 2006) was used to search these four genomes against the final assembly, and purge small contigs (<5,000 bp) that contained sequence matches (285 short contigs excluded). For larger scaffolds that contained possible embedded contaminated sequences, the high-quality ONT reads were mapped using Minimap2 over the regions. For those contaminated sites that had Nanopore reads spanning the contaminated region, the sequences were masked, and for those lacking nanopore support, the scaffold was split and/or trimmed to remove the contaminating sequence (seq 4 trimmed, seq 12 and 31 split into chromosome and unplaced scaffold). Finally, gaps of unknown size were standardised to 100 bp, and mitochondrial genome insertions into the nuclear genome were assessed using NUMTfinder (https://github.com/slimsuite/numtfinder) (Edwards et al., 2021) (none located). This paper primarily analyses S. vulgaris vAU1.0 (which we refer to has *S. vulgaris* vAU), while the final NCBI release (accession = JAGFZL000000000) is explicitly referred to as *S. vulgaris* vAU1.1 when relevant.

### 2.8 BUSCO versus BUSCOMP performance benchmarking

BUSCO-containing scaffolds from the Diploidocus primary haploid Supernova assembly of *Sturnus vulgaris* vAU were extracted into a reduced genome ‘pribusco’ assembly for additional BUSCO and BUSCOMP benchmarking (Supplementary Materials: Fig. S3). BUSCO v3.0.2b (Simão et al., 2015) (HMMER v3.2.1 (Wheeler & Eddy 2013), AUGUSTUS v3.3.2 (Stanke & Morgenstern 2005), BLAST+ v2.2.31 blast(Camacho et al., 2009), EMBOSS v6.6.0 (Rice et al., 2000)) was run in genome mode with the aves_odb9 dataset (n=4915) on: the non-redundant pseudodiploid (‘dipnr’), primary (‘pri’) and alternative (‘alt’) assemblies; BUSCO-containing scaffolds from the primary assembly (‘pribusco’); a reverse-complemented copy (‘revcomp’), combined with ‘pribusco’ to make a 100% duplicate assembly (‘duplicate’); a direct copy (‘copy’) combined with ‘duplicate’ to make a triplicated assembly (‘triplicate’); three randomly shuffled versions of ‘pribusco’ (‘shuffle1’, ‘shuffle2’, ‘shuffle3’), added in combination to ‘pribusco’ to generate datasets of increasing assembly size without increasing duplication levels (‘2n’, ‘3n’ and ‘4n’); ten straight repeats of the ‘pribusco’ run (‘rep0’ to ‘rep9’). All BUSCO results were processed with BUSCOMP v0.11.0 (Minimap2 v2.17). In addition to the full BUSCOMP analysis of all runs, the following subsets were grouped for analysis (Supplementary File 1, BUSCO v3 BUSCOMP output):

- Pseudodip: ‘dipnr’, ‘pri’ and ‘alt’. (Haploid versus diploid assemblies.)
- Core: ‘dipnr’, ‘pri’, ‘alt’, ‘pribusco’ and ‘revcomp’. (Assembly filtering and manipulation.)
- Duplication: ‘copy’, ‘duplicate’, ‘triplicate’. (Duplicating scaffolds.)
- Size: ‘shuffle1’, ‘shuffle2’, ‘shuffle3’, ‘2n’, ‘3n’, ‘4n’. (Increasing assembly size without duplication.)
- Replicates: ‘rep0’ to ‘rep9’.

The same analysis was repeated with BUSCO v5.0.0 (Simão et al., 2015) (SEPP v4.3.10 (Mirarab et al., 2012), BLAST v2.11.0 (Camacho et al., 2009), HMMer v3.3 (Wheeler & Eddy 2013), AUGUSTUS v3.3.2 (Stanke & Morgenstern 2005), Prodigal v2.6.3 (Hyatt et al., 2010), metaeuk v20200908 (Levy Karin et al., 2020)) and the aves_odb10 dataset (n=8338).

## 3. Results

### 3.1 *Sturnus vulgaris* vAU genome assembly

Genome assembly of *Sturnus vulgaris* vAU combined three different sequencing technologies for *de novo* genome assembly (10x Genomics linked reads, ONT long reads, and PacBio Iso-Seq full length transcripts) (Table 1), before a predicted reference-based scaffolding to the chromosome level using the high-quality reference assembly of *T. guttata* (NCBI REF: GCF_008822105.2). Approximately 109 Gb (97x coverage) of 10x linked read data (subsampled during assembly to 56x based on the estimated genome size of 1.119 Gb, barcode subsampling of 80%) were assembled with Supernova (v2.1.1) (Weisenfeld et al., 2017) (step 1) and converted to a primary haploid assembly (step2). We generated approximately 8 Gb of raw genomic reads using an ONT minion, which were reduced to 5 Gb after stringent filtering (Table 1). These data were used to scaffold the genome (step 3) and gap-fill (step 4), reducing the total number of scaffolds from 18,439 to 7,856, increasing the scaffold N50 from 1.76 Mb to 7.12 Mb, and decreasing the scaffold L50 from 146 to 39 (Supplementary Materials: Fig. S4). These measures were further improved after Iso-Seq scaffolding (step 5) (7,776 scaffolds, N50 7.12 Mb, and L50 38), followed by Pilon polishing using 10x linked reads (step 6). Finally, following haplotig removal (step 7), chromosomal alignment against the *T. guttata* reference genome (step 8) reduced the final number of scaffolds to 1,628 (N50 72.5 Mb, and L50 5) (Supplementary Materials: Fig. S4), with 98.6% of the assembly assigned to 32 putative nuclear chromosome scaffolds. While no whole mitochondrial genome insertions were found, 27 smaller mitochondrial pseudogenes (NUMTs) were located in *S. vulgaris* vAU1.1, with scaffold 31 (corresponding to the Z chromosome) containing the highest amount (Table S2).

**Table 1:** **Summary of sequencing data** for *Sturnus vulgaris* vAU genome assembly and annotation

| Genetic Data | Platform | Library | Library length/Mean Insert Size (kb) | Mean raw Read Length (bp) | Number of Reads | Number of bases (Gb) |
| --- | --- | --- | --- | --- | --- | --- |
| gDNA | Hiseq X Ten | Paired-end 10x Chromium | 51.7kb | 150 | 361,950,449 | 108.58 |
| gDNA | ONT MinION | Ligation | 47kb | 6,417 | 1,225,865 | 7.865 |
| cDNA | PacBio | Iso-Seq | Full transcripts (brain) (2.6 kb) | 12,000 | 20,558,110 | 38.650 |
| cDNA | PacBio | Iso-Seq | Full transcripts (heart + testes) (2.0 kb) | 10,000 | 18,985,944 | 29.496 |

#### Improvements to genome assembly completeness during scaffolding

Sequential steps of scaffolding, polishing, and quality control (Fig. 1, Supplementary Materials: Fig. S2, Table S3) improved the genome assembly statistics considerably from the initial Supernova *S. vulgaris* assembly (Supplementary Materials: Fig. S4). BUSCO completeness was approximately 94.6%, which was largely achieved by the initial assembly (92.9%), but somewhat improved over the additional assembly steps (Fig. 2a). The final BUSCO completeness score is comparable to other chromosome-level passerine assemblies on NCBI (Fig. 3a). BUSCO predictions are susceptible to base calling errors and can also fluctuate due to changes elsewhere in the genome assembly (Edwards 2019) (see section 2.8 BUSCO versus BUSCOMP performance benchmarking). As a consequence, BUSCO can under-report the true number of complete BUSCO genes in an assembly (Edwards et al., 2018; Field et al., 2020; Edwards et al., 2021). We therefore used BUSCOMP to compile complete BUSCO genes from across all stages of the assembly. Only 70 (1.4%) of the 4,915 Aves BUSCO genes were found to be “Missing” from all assembly versions, with 4,764 (96.9%) rated “Complete” in at least one stage (Fig 3a, BUSCOMP).

**Figure 2:**
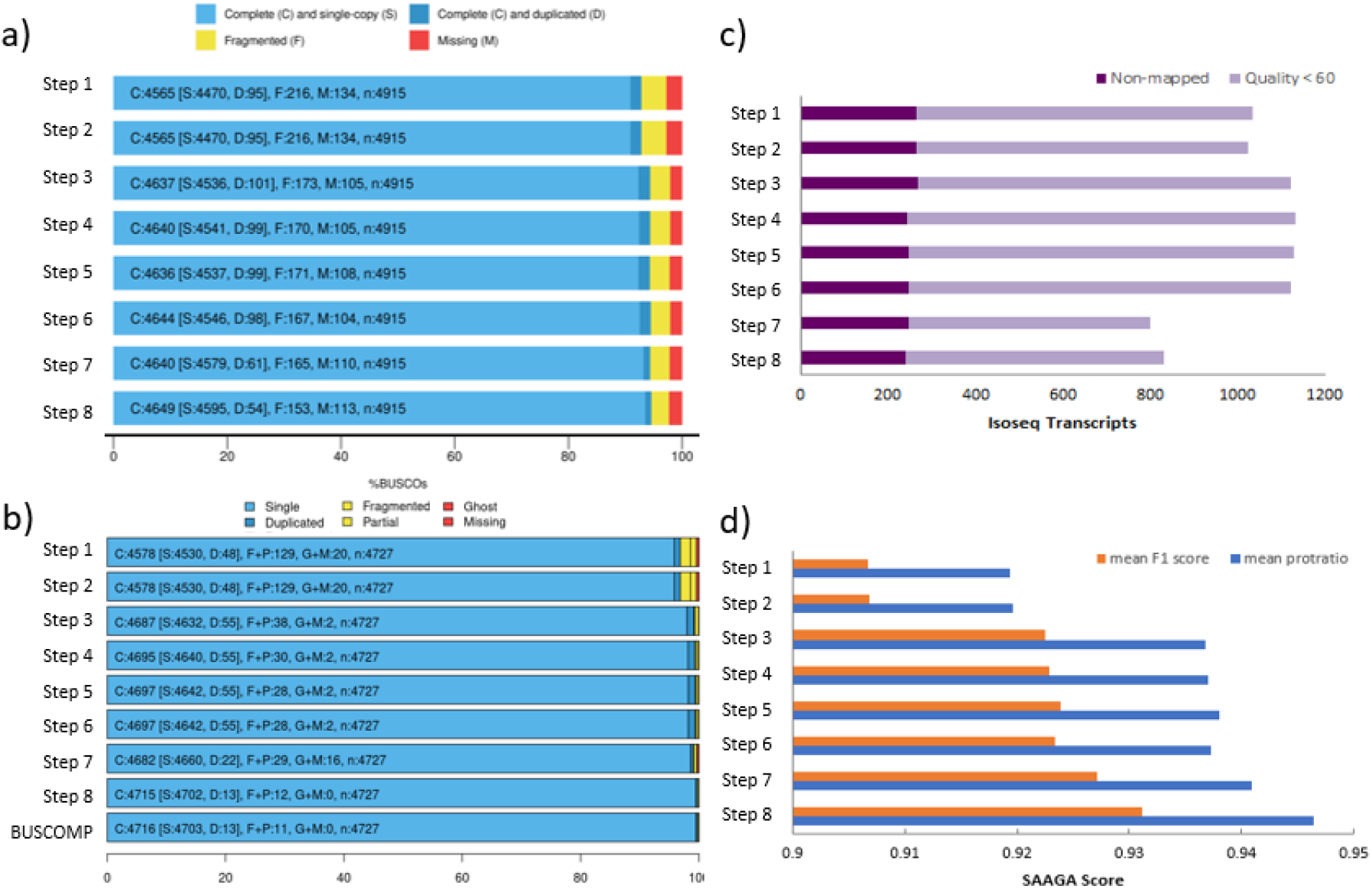
*Sturnus vulgaris* vAU assembly steps overview. Quality and completeness assessments for eight sequential assembly steps: step 1 (Supernova assembly), step 2 (Diplodocus primary assembly), step 3 (Sspace -Longreads scaffolding), step 4 (Gapfinisher gapfilling), step 5 (L_RNA_scaffolder), step 6 (Pilon polishing), step 7 (Diplodocus clean up), and step 8 (Satsuma2 Chromosome scaffolding). **a)** BUSCO (Aves, n=4,915) completeness rating summaries for the sequential steps of *S. vulgaris* genome assembly. **b)** BUSCOMP completeness results for the 4,727 BUSCO genes identified as single copy and complete in one or more assembly stages. The final BUSCOMP row compiles the best rating for each gene across all eight steps. **c)** The number of Iso-Seq reads that failed to map to each assembly step. **d)** SAAGA annotation scores of mean protein length ratio (blue) and F1 score (orange) (see Methods for details).

**Figure 3:**
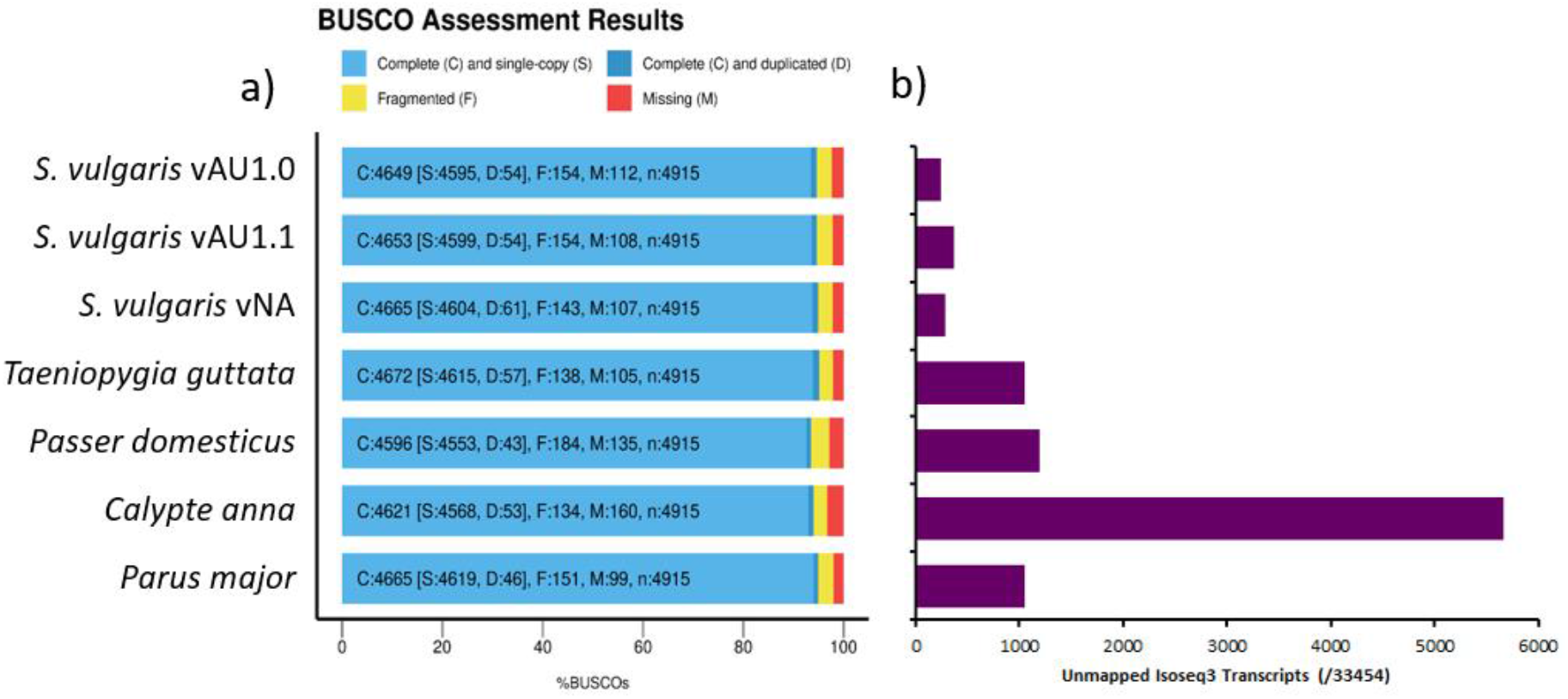
Assessment of *Sturnus vulgaris* and comparison avian reference assemblies. **a)** BUSCO (Aves) assessments of assembly completeness of *S. vulgaris* vAU1.0, and the NCBI uploaded genome *S. vulgaris* vAU1.1, presented alongside *S. vulgaris* vNA and four recent high-quality avian reference genomes (*Taeniopygia guttata* assembly accession GCF_008822105.2, *Passer domesticus* assembly accession GCA_001700915.1, *Calypte anna* assembly accession GCA_003957555.2, *Parus major* assembly accession GCA_001522545.3). **b)** Total number of Iso-Seq transcripts that failed to map to each assembly.

The final assembly had the fewest unmapped Iso-Seq reads (Fig. 2b), with the largest improvement seen post gap-filling, followed by chromosome scaffolding. An increase in missing Iso-Seq transcripts was observed after scaffolding with the Iso-Seq reads themselves, and post long-read scaffolding, due to reads no longer partially matching at scaffold ends. Polishing caused a minimal improvement on the total number of mapped Iso-Seq reads, and none were lost during scaffold clean-up with Diploidocus (runmode purgehaplotig and vecscreen). Assessment using GeMoMa and SAAGA revealed that across these assembly steps we see a generally consistent increase in the quality of the predicted proteins during annotation (Fig. 2c), with the largest increases occurring post long-read scaffolding, followed by chromosome scaffolding, and then scaffold clean-up.

Of the 33,454 high-quality isoform transcripts in the PacBio Iso-Seq data, only 241 failed to map to the final genome assembly, a 17.2% decrease compared to the 291 that failed to map to *S. vulgaris* vNA (Fig. 3b).

#### Final genome assembly size, heterozygosity, and contiguity

The *S. vulgaris* vAU assembly of 1,049,838,585 bp covers approximately 93.78% of the total estimated 1.119 Gb genome size (Supplementary Materials: Appendix 2 Validation of Supernova genome size prediction using Jellyfish). A similar estimation of genome completeness was reported by K-mer Analysis Toolkit (kat), with the raw read1s (forward reads) estimating a genome completeness of 96.7% (estimated genome size 1.125 Gb, estimated heterozygosity rate 0.57%) and read2s (reverse reads) estimating a genome completeness of 95.92% (estimated genome size 1.135 Gb, estimated heterozygosity rate 0.54%) (Supplementary Materials: Fig. S5). Predicted genome sizes based on either read1s or read2s using kat were slightly larger than the estimation generated by Jellyfish using all the read data, however the length range was relatively consistent (1.119-1.135 Gb). This assembly reports a scaffold N50 of 72.5 Mb and L50 of 5, with a total of 1,628 scaffolds (Table 2); 98.6% (1,035,260,756 bp) of the sequence length has been assigned to the 32 putative nuclear chromosomes (identified via the *T. guttata* v3.2.4 assembly), plus a mitochondrial genome. The final assembly contains 14 macrochromosomes (> 20 Mb, as described in Backström et al., 2010), with relative sizes appearing in consensus with known karyotype of *S. vulgaris* (Calafati & Capanna 1981). Macrochromosome scaffolds account for 81.9% of the total assembly size, with the remainder on microchromosomes (16.9%) or unplaced scaffolds. While these large scaffolds remain only putative chromosomes assuming karyotype orthology until they can be validated with further read data, increased completeness scores post chromosomal alignment across all assembly assessment metrics (Fig. 2) support the assembly structure.

**Table 2:** *Sturnus vulgaris* overview of assembly statistics for vAU1.0, vAU1.1, and vNA, assessed using BUSCOMP.

|  | *Sturnus vulgaris*<br>vAU1.0 | *Sturnus vulgaris*<br>vAU1.1 | *Sturnus vulgaris*<br>vNA |
| --- | --- | --- | --- |
| Total length (bp) | 1,049,838,585 | 1,043,825,671 | 1,036,755,994 |
| Number of scaffolds | 1,628 | 1,344 | 2,361 |
| Scaffold N50 (bp) | 72,525,610 | 72,244,370 | 3,416,708 |
| Scaffold L50 | 5 | 5 | 89 |
| Largest scaffold (bp) | 151,927,750 | 151,503,485 | 11,828,398 |
| Mean scaffold length (bp) | 644,864.0 | 776,656.01 | 439,117.3 |
| Median scaffold length (bp) | 1,337 | 1,343 | 4,856 |
| Number of Contigs | 23,815 | 23,340 | 22,666 |
| Contig N50 (bp) | 145,864 | 147,322 | 147,183 |
| Contig L50 | 2,030 | 2,010 | 1,908 |
| Gap (N) length (bp) | 13,242,113 (1.26%) | 0.74% | 23,939,528 (2.31%) |
| GC (Guanine-Cytosine) content (%) | 41.73% | 41.72% | 41.49% |

### 3.2 *Sturnus vulgaris* vAU whole transcriptome data analysis

We generated approximately 68 Gb of PacBio Iso-Seq whole transcript (39,544,054 subreads) (Table 1). This produced a total of 33,454 clustered high-quality (predicted accuracy ≥ 0.99) reads, and 157 clustered low-quality (predicted accuracy < 0.99) reads (Supplementary Materials: Table S4). These high-quality read data were used to improve the scaffold assembly of the genome using l_rna_scaffolder (see section 2.1) and assess genome completeness (using count comparison of unmapped Iso-Seq reads, see section 2.5.2). After being passed through the Tama *collapse* pipeline, a total of 28,448 non-redundant transcripts were retained to create the final *S. vulgaris* vAU transcriptome, which was used for gene prediction when completing the annotation of the genome assembly. This final three tissue (brain, gonad, heart) Iso-Seq transcriptome had a moderate level overall BUSCO completeness of around 63% that compares to other avian Iso-Seq transcriptomes (Fig. 4a), with a wide range of gene ontology terms identified in the final Iso-Seq transcript list (Fig. 4b) that resembled other avian Iso-Seq GO term distributions (Yin et al., 2019).

**Figure 4:**
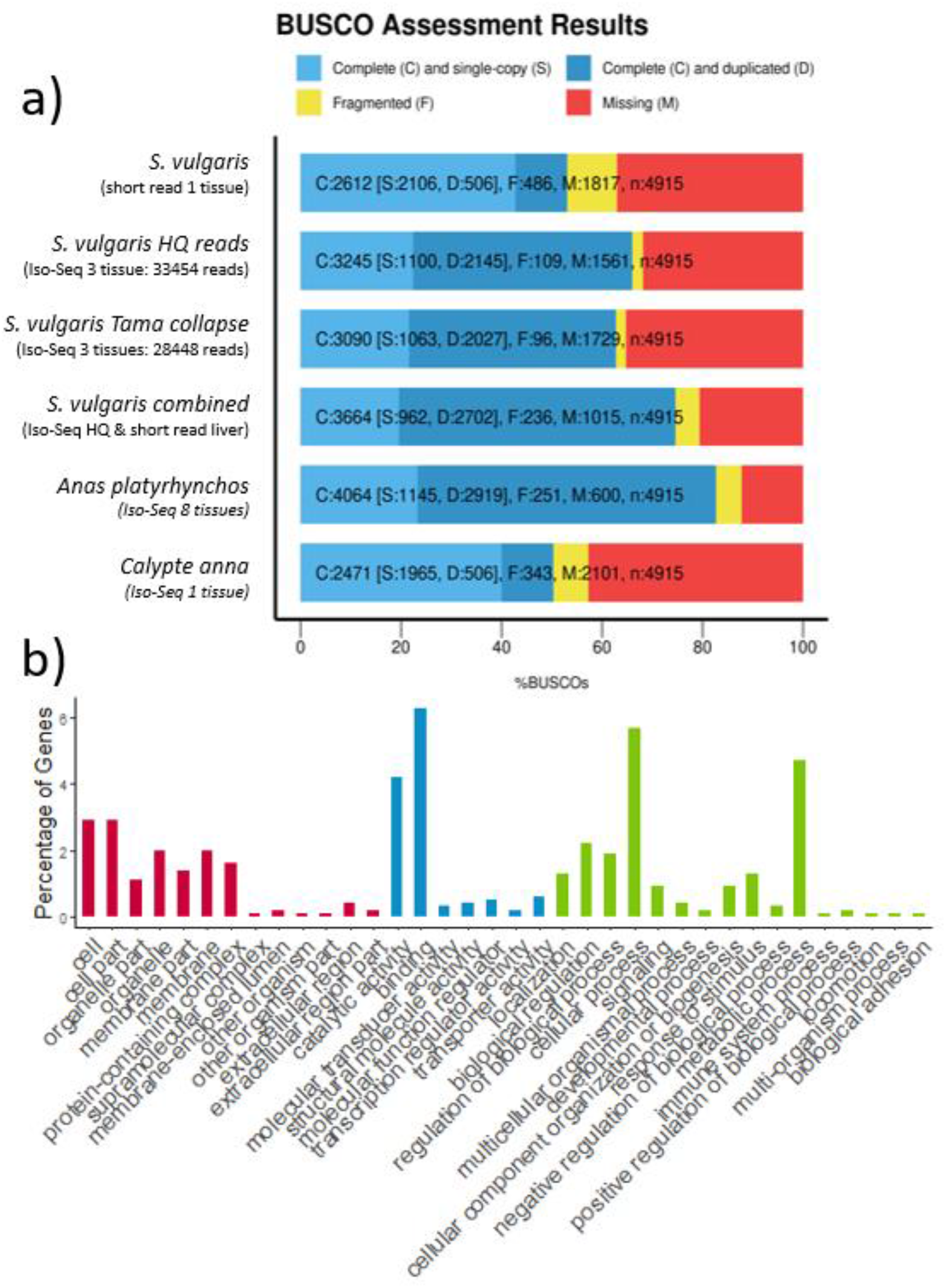
Assessment of 3 tissue Iso-Seq (brain, gonad, heart) *Sturnus vulgaris* transcriptome. **a)** BUSCO (aves) rating summaries for *S. vulgaris* short read liver transcriptome, the high-quality Iso-Seq *S. vulgaris* transcript produced though the Iso-Seq v3.3 pipeline, the final *S. vulgaris* transcriptome produced by Tama *collapse* pipeline, and combined high-quality Iso-Seq and short read liver transcripts, alongside two other avian Iso-Seq transcriptomes (*Anas platyrhynchos* using pectoralis, heart, uterus, ovary, testis, hypothalamus, pituitary and 13 days-old embryo tissue (Yin et al., 2019), and *Calypte anna* using liver tissue (Workman et al., 2018)). **b)** Breakdown of major GO terms in the sequenced Iso-Seq reads, with Cellular Component (red) Molecular Function (blue) and Biological Process (green).

### 3.3 *Sturnus vulgaris* genome annotation

The initial annotation produced by GeMoMa, informed by the 26 avian genome annotations available at the time on Ensembl (Supplementary Materials, Table S1), predicted 21,539 protein coding genes, with 97.2% BUSCO completeness (93.1% complete when longest protein-per-gene extracted with SAAGA) (Fig. 5). The initial Maker2 annotation reported 13,495 genes, and a BUSCO completeness of 79.5% (Fig. 5). The merged final annotation reported a BUSCO completeness of 98.2% (Fig. 5a), and this annotation predicted a total of 21,863 protein-coding genes and 79,359 mRNAs. There was an average of 10.7 exons and 9.7 introns per mRNA, with an average intron length of 3,364 (Table 3). Of these, 1,764 are single-exon genes and 2,330 single-exon mRNA. Predicted coding sequences make up 5.4% of the assembly, with 44.77% remaining outside any gene annotation (Fig. 5b).

**Table 3:**
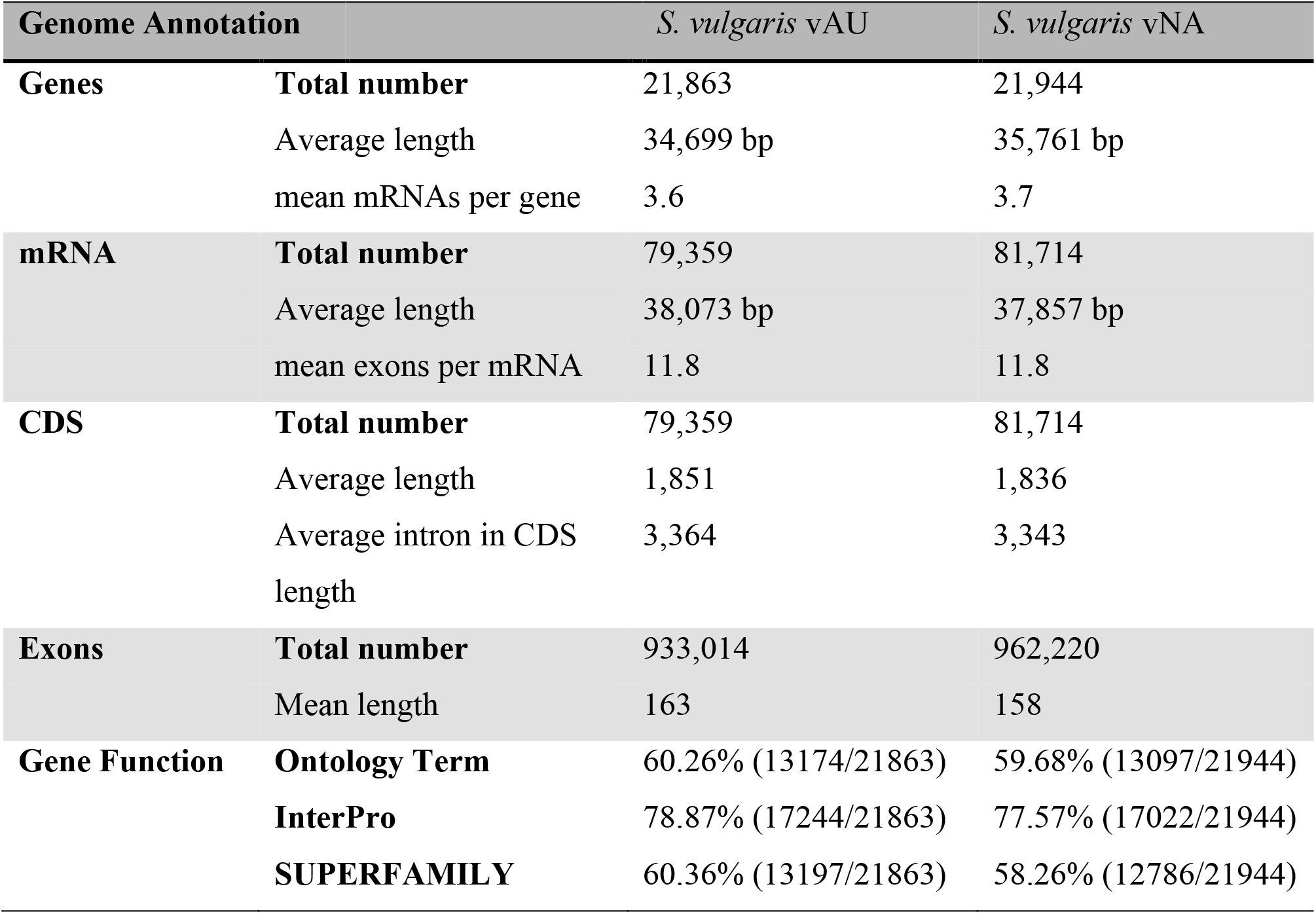
Summary of genome annotation of *Sturnus vulgaris* vAU and vNA assemblies. Statistics extracted using AGAT *agat_sp_functional_statistics.pl*.

**Figure 5:**
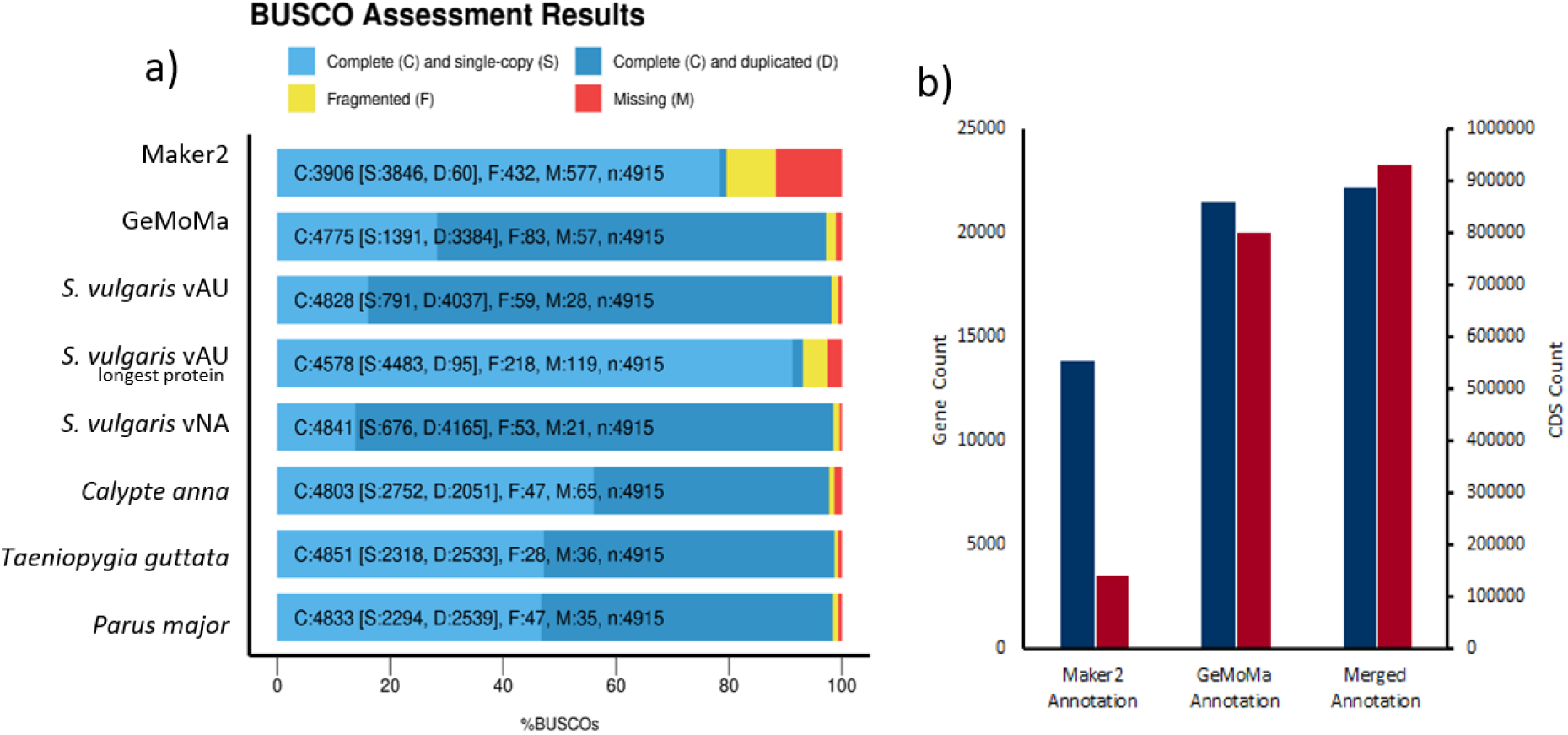
*Sturnus vulgaris* assessment of annotation. **a)** BUSCO (Aves) assessments of initial Maker2 and GeMoMa assemblies, the final *S. vulgaris* vAU annotation, the final annotation with the longest protein-per-gene extracted using SAAGA, the final *S. vulgaris* vNA annotation (combined GeMoMa and Maker2 annotation), and the ensemble annotations of three additional passerines. **b)** The number of genes (blue) and CDS (red) in the Maker2 annotation, GeMoMa annotation, and merged annotation.

The predicted transcripts were mapped using SAAGA to the Swiss-Prot database, with 66,890 transcripts returning successful hits (84.3%) and 12,469 transcripts remaining unknown (15.7%) for the final annotation (Fig. 6a). The known proteins had an average length of 652 amino acids (aa) and the unknown proteins had an average length of 426 aa (Fig. 6a). Most of the predicted proteins were of high quality, with around 56% of them having an F1 score (see Methods) of greater than 0.95 (Fig. 6b). Similar results were seen when the *Gallus gallus* reference proteome was used, with 69,714 known proteins of average length of 646 aa, 9,645 known proteins of average length of 401 aa, and the final merged annotation having the same F1 score distribution (Fig 6c & 6d).

**Figure 6:**
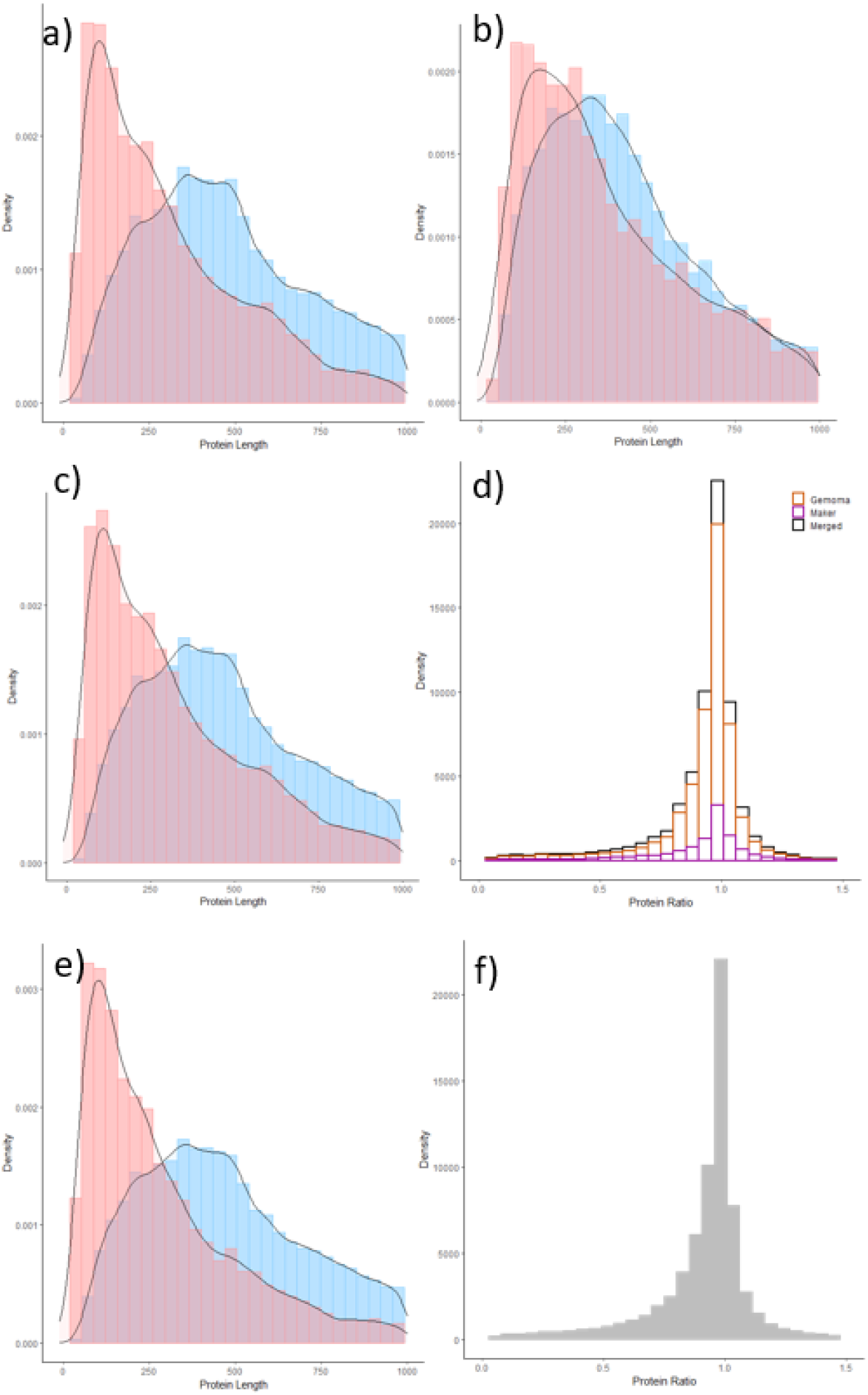
Summary of predicted annotated proteins. **a)** Protein lengths for known proteins (blue, with a located Swiss-Prot comparison) and unknown proteins (red, those that did not map to Swiss-Prot) for the GeMoMa annotation compared to Swiss-Prot. **b)** Protein lengths of known and unknown proteins for the Maker2 annotation compared to Swiss-Prot. **c)** Protein lengths of known and unknown proteins for the merged GeMoMa and Maker2 annotation compared to Swiss-Prot. **d)** Protein length ratio between output from SAAGA for all known Swiss-Prot proteins (where a score close to 1 indicates a high-quality gene annotation, protein length ratio calculated as annotated protein length / best Swiss-Prot reference protein length) (merged annotation = black, GeMoMa annotation = orange, Marker2 annotation = purple). **e)** Protein lengths of known and unknown proteins for the merged GeMoMa and Maker2 annotation compared to *Gallus gallus* reference proteome (UP000000539_9031). **f)** Protein length ratio between output from SAAGA for the merged annotation against the *Gallus gallus* reference proteome.

The GeMoMa annotation had similar protein quality patterns, with 57,026 known proteins (average length 664 aa), and 10,400 unknown proteins (average length 401 aa) (Fig. 6e). The Maker2 displayed much greater similarity in protein length histogram between known and unknown proteins, with shorter proteins with known homologs (average length 565 aa), but longer unknown proteins (average length 549 aa) (Fig. 6f). The *S. vulgaris* vNA annotation final merged annotation had extremely similar statistics to the final *S. vulgaris* vAU annotation, with an average known protein length of 650 aa, and an average unknown protein length of 407 aa (Supplementary Materials: Figure S2b).

### 3.4 *Sturnus vulgaris* genome-wide patterns of genomics features

Transcript density compared between mapped Iso-Seq reads and predicted transcripts in the final annotation displayed similar patterns, with some minor variation in patterns between the two tracks (Fig. 7; track 1). Final predicted gene densities (Fig. 7; track 2) were largely following the patterns seen in transcript densities. Further, patterns of transcript and gene numbers across the genome track relatively consistently to GC content (Fig. 7; track 4).

**Figure 7:**
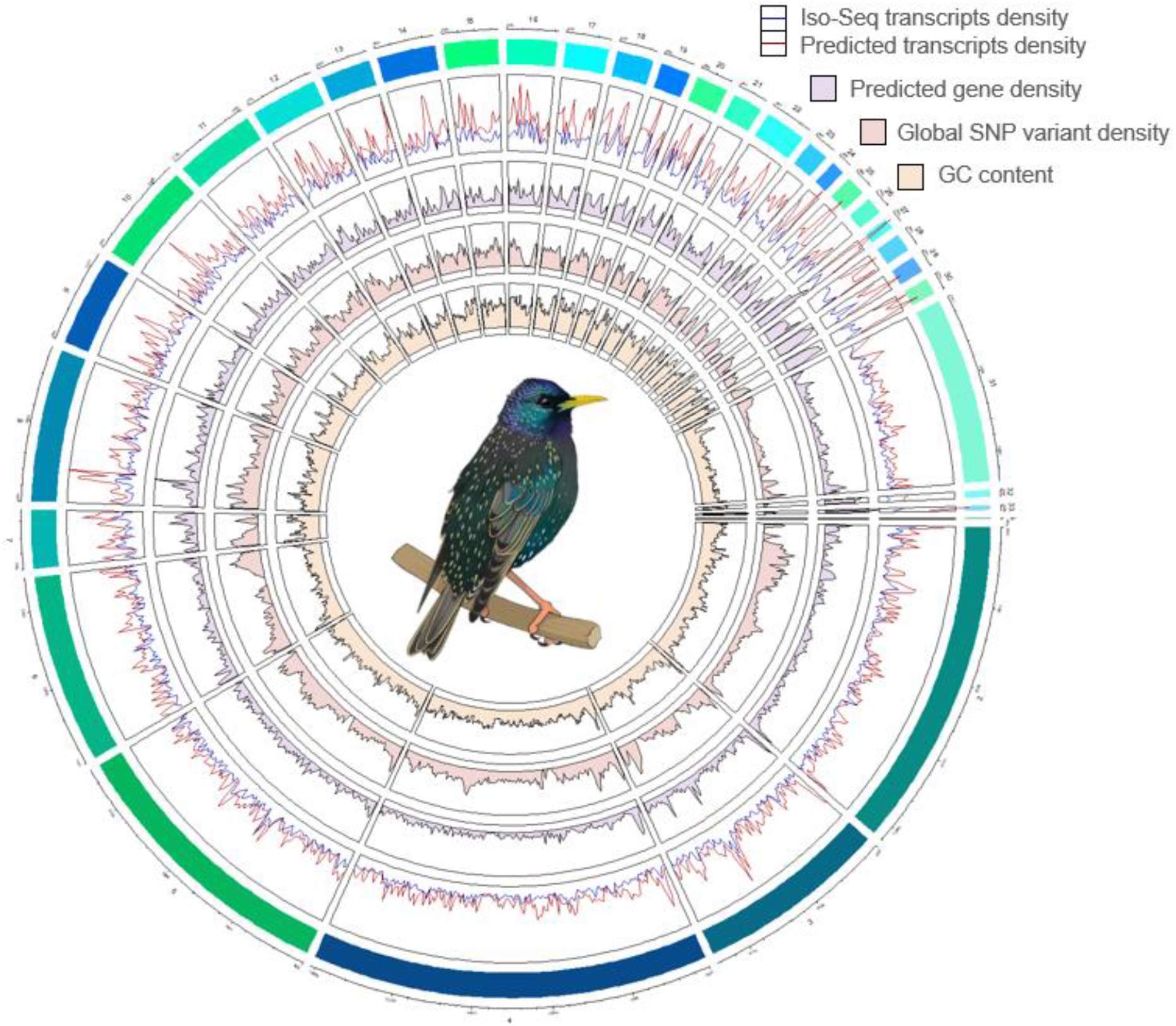
CIRCLIZE plot of the 33 main chromosomal scaffolds. (32 putative autosomes plus mtDNA) in the Sturnus vulgaris (*S. vulgaris* vAU) genome assembly (>98% of the total assembly length). The tracks denote variable values in 1,000,000 bp sliding windows. From the outermost track in, the variables displayed are track 1 (Iso-Seq transcripts as blue line, final annotation transcripts as red line), track 2 (final annotation gene counts, purple area), track 3 (variant density, red area), and track 4 GC content (yellow area).

Global whole genome variant data (Fig. 7; track 3) revealed genomic regions where variant density is low or non-existent, indicative of high genetic conservation across the species, and genomic regions where variant density peaks are indicative of variant hotspots. Interestingly, we see regions of high conservation corresponding to peaks in gene and/or transcript numbers (e.g., midway through chromosome 4), which may be indicative of regions of highly conserved genes and possibly centromere locations.

### 3.5 BUSCO versus BUSCOMP performance benchmarking

For the non-redundant pseudodiploid *S. vulgaris* vAU Supernova assembly, BUSCOMP revealed differences in the BUSCO ratings of scaffolds dependent on the assembly background (Fig. 8). Despite the primary (‘pri’) assembly being a subset of the non-redundant pseudodiploid (‘dinpnr’) assembly, it identified more “Complete” BUSCO genes (4,565 versus 4,532) with fewer “Missing” (131 versus 171) (Fig. 8a). The alternative assembly (‘alt’) subset similarly returned a partially overlapping set of BUSCO genes with ‘dipnr’, including some not found in ‘dipnr’ or ‘pri’: in total, only 101 genes were missing from all three assemblies. Reducing the primary assembly to the 968 (of 18,439) scaffolds containing a complete BUSCO gene (‘pribusco’), increased the number of complete genes from 4,565 to 4,586 and reduced the number missing from 131 to 112. Most unexpectedly, reverse complementing these scaffolds reduced the number of BUSCO genes rated “Complete” by two, and increased the number “Missing” by fifteen (Fig. 8a). All five assemblies returned complete BUSCO genes that were fragmented or missing in all the other four assemblies (Supplementary File 1, BUSCOMP v3 results), for a combined total of 4,760 complete and only 74 missing.

**Figure 8.**
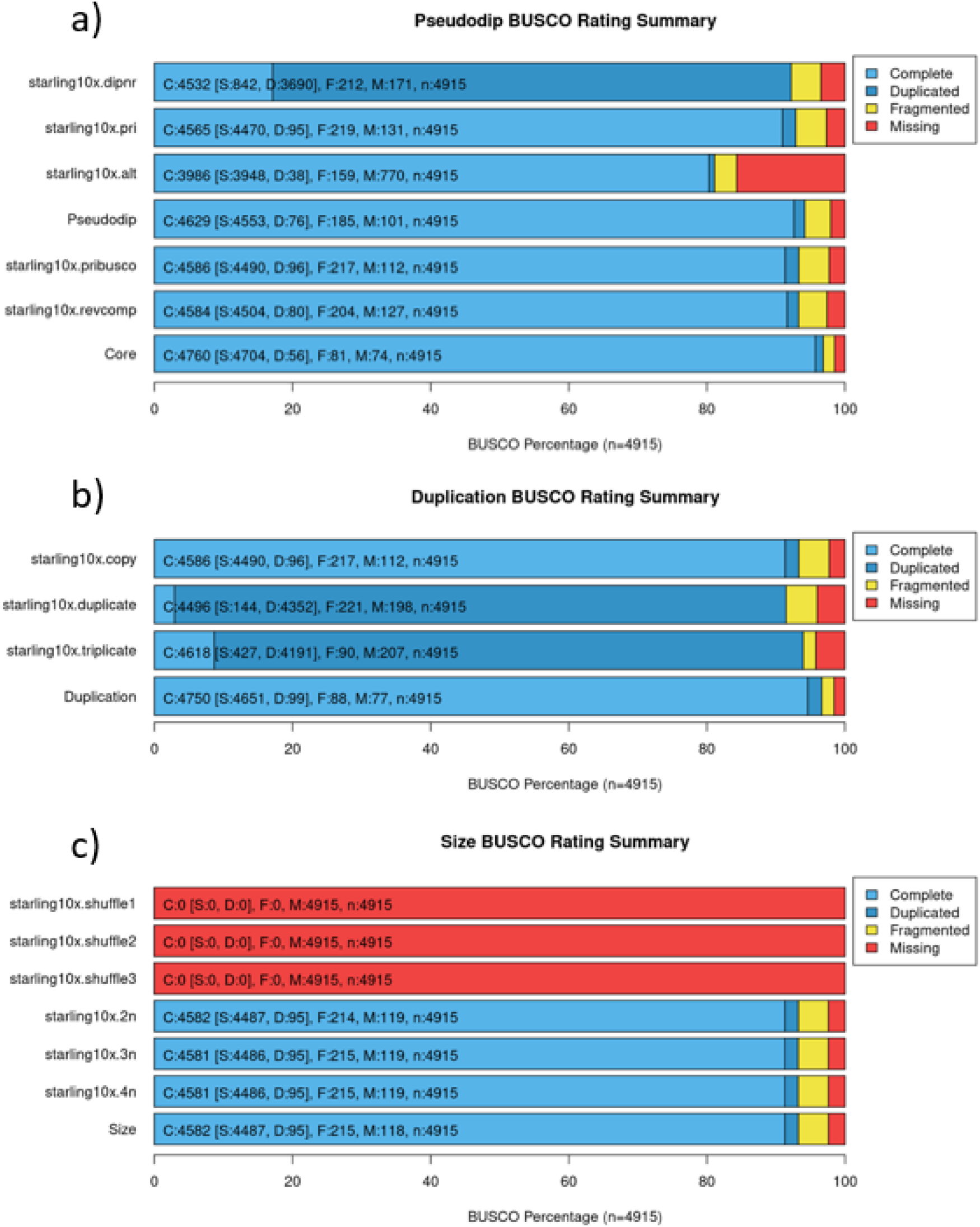
Compiled BUSCO results for benchmarking data.

Adding direct or reverse-complemented copies of the ‘pribusco’ scaffolds increased the number of “Duplicated” genes, but still returned single copy complete genes (Fig. 8b). Doubling and then tripling the assembly size also increased the number of “Missing” genes from 112 to 198 (‘duplicate’) and then 207 (‘triplicate’). As before, these summary numbers hide some gene gains as well as gene losses; only 77 genes are missing from all three BUSCO runs, with 4,750 returned as complete by at least one. Adding randomly shuffled versions of the ‘pribusco’ scaffolds only had a marginal effect on BUSCO ratings, with four (‘2n’) to five (‘3n’, ‘4n’) fewer complete genes returned and seven additional genes missing following addition of the random sequences (Fig. 8c). Ten replicate analyses of the ‘pribusco’ scaffolds returned identical results (Supplementary File 1, BUSCOMP v3 results).

In contrast, BUSCOMP completeness is much more consistent across all datasets, with the primary assembly returning the same numbers of complete, partial/fragmented and missing genes as the pseudodiploid assembly (Supplementary File 1, BUSCOMP v3 results). Similarly, reverse complementing scaffolds or increasing genome size gives no difference to the completion statistics. Unlike BUSCO, BUSCOMP rates 100% of complete BUSCO genes for duplicated or triplicated scaffolds as ‘Duplicate’ rather than ‘Single Copy’. Most reassuringly, every complete BUSCO gene returned by a variant or subset of the pseudodiploid assembly is also returned as ‘Complete’ in the pseudodiploid assembly itself. Results using BUSCO v5 and the updated lineage data were qualitatively the same as v3, showing largely identical trends (Supplementary File 2, BUSCOMP v5 results). The exception is that reverse-complementing scaffolds reduced the complete BUSCO genes by one (7,555 to 7,554) and increased the number missing by one (391 to 392). Curiously, this was not reflected by analysis of the duplicated scaffolds, in which all 7,555 ‘pribusco’ complete genes were returned as complete and duplicated. It should be noted that the ‘pribusco’ scaffolds for the v5 analysis are missing a greater proportion of the BUSCOMP-compiled single copy complete BUSCO genes because they were still defined from v3 data.

## 4. Discussion

Here, we present a high-quality, near-complete reference genome for the European starling, *Sturnus vulgaris* vAU, with chromosome-level scaffolding that assigns 98.6% of the genome assembly length to 32 putative nuclear chromosome scaffolds. We demonstrate the utility of both transcripts and gene annotation in validating *S. vulgaris* vAU assembly processes. BUSCOMP, Iso-Seq transcript, and SAAGA annotation assessment were largely in agreement with one another, though each provided additional fine-scale feedback on assembly improvements achieved by each assembly step. These analyses highlight the benefits of these complementary assessment approaches in ensuring that aspects of genome quality are not sacrificed to improve non-specific assembly quality metrics, such as N50. We also present a second, North American, genome assembly, *S. vulgaris* vNA (GCF_001447265.1). Overall, the *S. vulgaris* vAU assembly improved genome assembly statistics over the *S. vulgaris* vNA genome, with a greater percentage of the estimated 1.119 Gb genome represented (94% vs 93%), an increase of scaffold N50 from 3.42 Mb to 72.5 Mb, and a decrease in scaffold L50 from 89 to 5. The *S. vulgaris* vNA still has good assembly statistics (Table 2, Table S3) and has a marginally higher BUSCO completeness (Fig. 3a) and BUSCOMP completeness (Supplementary Materials: Fig. S6) of approximately 20 BUSCO sequences. There is increasing recognition of the importance of pan-genomes (genome assemblies that differentiate between genes/regions shared by all members of the species, and dispensable or rare genes/regions) (Hirsch et al., 2014; Sherman & Salzberg 2020), which are essential for many model organisms (Vernikos et al., 2015). Having these two high-quality *de novo* assemblies from different populations will improve future genomic work on the global invasive populations of this species, and facilitate review of structural variation (e.g., inversions) that may exist across different populations. It should be noted, however, that the final scaffolding step for *S. vulgaris* vAU assumed structural conservation between the starling and zebra finch and thus future synteny analyses may want to use the earlier assembly step.

### 4.1 BUSCO and BUSCOMP assembly completeness assessment

BUSCO (Simão et al., 2015) is an extremely useful and widely-used used assembly assessment tool, providing information on which conserved lineage specific genes are present, fragmented, or absent from a genome assembly. The program, however, can suffer from inconsistent BUSCO gene identification, where a particularly BUSCO may be dropped from a report due to changes elsewhere in the assembly (Edwards 2019), which can result in under-reporting of assembly completeness (Edwards et al., 2018; Field et al., 2020; Edwards et al., 2021). Here, we confirm this behaviour on benchmarking datasets derived from the *S. vulgaris* vAU pseudodiploid 10x linked read assembly (Supplementary Materials: Fig. S3, 8). Adding and removing scaffolds can both alter the BUSCO ratings for “Complete” genes within the unchanged scaffolds (Fig. 8, Supplementary File 1, BUSCOMP v3 results, Supplementary File 2, BUSCOMP v5 results). Many of these changes are likely to be the consequence of changes in score thresholds and/or gene prediction models. However, we also demonstrate some unexpected behaviours that are harder to explain, such as changes to BUSCO gene ratings when scaffolds are reverse complemented (Fig 8a).

This unpredictable variability in the identification of BUSCOs across genome assembly versions poses some obvious challenges when trying to compare alternate versions of the same assembly. This is particularly true when trying to interpret small changes in BUSCO ratings as assemblies near completion. In addition, an important feature of BUSCO is that it incorporates sequence quality in the context of the gene prediction models it generates. This is desirable for assessing final assembly quality, but can present problems when comparing early assembly stages, prior to error-correction by “polishing”. BUSCOMP (https://github.com/slimsuite/buscomp) is robust to differences in assembly size, base-calling quality, and rates the “completeness potential” of an assembly based on the presence of genes first identified for that species by BUSCO. Here, we used BUSCOMP analysis of sequential assembly steps to gain a more accurate understanding of how assembly decisions affected genome completeness (Fig. 2, Supplementary Materials: Fig. S4). BUSCOMP analysis can then be complemented by other tools, such as KAT (Mapleson et al., 2017), SAAGA (https://github.com/slimsuite/saaga), and BUSCO itself to get additional assessment of sequence quality.

### 4.1. Transcript- and annotation-guided *Sturnus vulgaris* vAU genome assembly

The assembly of the *S. vulgaris* vAU genome was improved by assessing mapped Iso-Seq whole transcripts and quality scores of predicted proteins from homology-based annotation. Mapping of the high-quality Iso-Seq reads proved to be an extremely fast method of assessment (33,454 Iso-seq sequences mapped in <5 mins with 16 CPU cores), while the GeMoMa and SAAGA compute time of 12 hrs per assembly was roughly comparable to BUSCO (approximately 50 CPU hrs per assembly on an average machine), though more computationally intensive (GeMoMa ran for approximately 200 CPU hours per assembly, and SAAGA ran for approximately 8 CPU hours per assembly). Over the eight sequential assembly steps, there was a decrease in unmapped Iso-Seq reads, indicating improved sequence representation, with gap-filling yielding the greatest change. Similarly, the quality of annotated proteins predicted by GeMoMa, as assessed by SAAGA, demonstrated ongoing improvements through ONT scaffolding, clean-up, and chromosome alignment. It is also noteworthy that increases in large-scale sequence connectivity using the *T. guttata* genome (Peona et al., 2018) improved the assembly’s performance across all metrics, including completeness estimates, although future Hi-C analysis will be required to confirm the predicted genome structure.

Further, BUSCOMP provided an important means of standardising BUSCO annotation ratings across the multiple assembly steps. This method, together with the mapped Iso-Seq reads, can deal with the unpolished intermediary genome steps, and does not suffer the same sequence identification accuracy issues as the traditional stand-alone BUSCO analysis. Together, the standardised assessment reported by BUSCOMP, and the comprehensive and genome/species specific set of genes provided by Iso-seq and GeMoMa/SAAGA showcase the complementary features of these annotation approaches for assembly assessment.

### 4.2. Improvements to contiguity and completeness during *Sturnus vulgaris* vAU genome assembly

Several alternative assembly pipelines were assessed (Supplementary Materials: Fig. S2), with upstream assembly decisions based primarily on establishing reasonable base assembly statistics (scaffold N50, scaffold L50, contig numbers). Assembly size increased during scaffolding steps, due to estimated bases in gaps, while a decrease in assembly size was only seen during scaffold clean up using *Purgehaplotigs*. Of all the scaffolding steps, scaffolding with the low coverage ONT long reads resulted in the greatest decrease of scaffold L50 (146 scaffold to 39 scaffold, Supplementary Materials: Fig. S4d) and total scaffold number (18,439 scaffolds to 7,856 scaffolds, Supplementary Materials: Fig. S4a). It has previously been shown that even low coverage of ONT data in conjunction with 10x may produce high-quality genome assemblies (Ma et al., 2019). This was true for our data, which demonstrates the utility of even low coverage, long read sequencing (approximately 4.5% coverage based on the estimated genome size of 1.119 Gb) in greatly improving the contiguity of scaffolds generated by short read genome assemblers (though Hi-C data may serve this purpose at a lower cost to scaffold ratio and may assist in identifying misassemblies, which is often not a focus of long-read scaffolding tools). While additional scaffolding using the Iso-Seq whole transcripts did not result in a large increase in continuity (Supplementary Materials: Fig. S4), the Iso-Seq reads were nevertheless were able to scaffold some sequences that failed low coverage ONT scaffolding, reducing the total scaffold count by approximately 100 (7,856 scaffold to 7,776 scaffolds, Supplementary Materials: Fig. S4a). This long-read transcript scaffolding served to minimise the number of fragmented genes in the final assembly, helping downstream analysis and gene prediction models. The final assembly maintained reasonably short contig N50 and high contig L50, which will only be improved with much more extensive long-read sequencing of the species. Nevertheless, scaffolding the *S. vulgaris* genome against that of *T. guttata* was able to further scaffold the genome to a predicted chromosome level, assigning 98.6% of the assembly to previously characterised chromosomes. In support of the assumed synteny of this step, we saw small increases in assembly quality and completeness metrics.

The final two assembly steps (contig clean up and chromosomal alignment) were primarily guided by high BUSCO scores and low missing Iso-Seq transcripts (Supplementary Materials: Figure S3). Diploidocus *vecscreen* did not flag any contamination and so did not result in any assembly decreases. Over-pruning of contigs during clean up (using Diploidocus *DipCycle* which is stricter than just *Purgehaplotigs*) resulted in too many (>1,000) discarded scaffolds that decreased assembly completeness scores, most likely because of low coverage ONT long read data. While this drastically improved assembly statistics, this came at the cost of dropped BUSCO sequences. Lastly, assembly duplication analysis using KAT agreed with BUSCO results, indicating there was little final assembly sequence duplication when comparing to raw read k-mer counts (Supplementary Materials: Fig. S5).

### 4.3. *Sturnus vulgaris* vAU transcriptome

When comparing the completeness of this new starling transcriptome data to existing Illumina short read transcript data produced using liver tissue (Richardson et al., 2017), we see an increase of about 20% in BUSCO completeness, with a particularly large increase in the number of duplicated BUSCO, a result of the alternate transcript isoforms captured through the Iso-Seq. Assessing the effect the Tama pipeline had on BUSCO completeness, we see a small drop in complete BUSCOs (Fig. 2a) that appear to have been lost during the mapping to genome assembly step. Finally, comparing our final transcriptome to two other avian Iso-Seq transcriptomes gives an indication of how much unique transcript information is added by the addition of tissues into pooled Iso-Seq sequencing runs. The single tissue Iso-Seq liver transcriptome of *Calypte anna* (Anna’s hummingbird) (Workman et al., 2018) reported similar BUSCO completeness to the short read *S. vulgaris* liver transcriptome. The eight tissue Iso-Seq transcriptome of *Anas platyrhynchos* (mallard) (Yin et al., 2019) yielded an increase of 30% in complete BUSCOs, consistent with the expectation that our three-tissue Iso-Seq library will be missing a number of tissue-specific genes.

### 4.4. *Sturnus vulgaris* genome annotation

Of the approximately 22,000 genes reported in the final annotation, 65% were from GeMoMa, and 35% from Maker2, with the source being randomly selected for common annotation. Maker2 predicted a higher number of genes in *S. vulgaris* vNA versus *S. vulgaris* vAU (15,150 vs 13,495), while GeMoMa predicted a higher number of genes in the *S. vulgaris* vAU genome (21,539 vs 20,414). The ratio in predicted Maker2 and GeMoMa was more biased towards the homology-based predictor, with an approximate ratio of 1:5 between Maker2 and GeMoMa (Fig. 5b). Merging of the MAKER2 annotation to the GeMoMa annotation resulted in an increase in 1.1.% in BUSCO completeness. Duplication levels were much higher in the GeMoMa annotation when compared to Maker2 (Fig. 5a). This is not unreasonable, as the GeMoMa annotation will be biased toward well-characterised genes and so may contain more transcripts per gene (Fig. 5b), whereas Maker2 will inform the prediction of more taxon or possibly species-specific coding sequences. High congruence between Iso-Seq and predicted transcript numbers indicate regions of accurate annotation predictions (Fig. 7). In contrast, Iso-Seq transcripts that are dissimilar or much lower to the predicted transcript densities, are either genomics regions producing tissue specific transcripts not captured by their brain, testes, or muscle, or more likely annotated transcript overprediction.

For the final *S. vulgaris* vAU annotation, the predicted proteins of unknown origin (those that failed to map to Swiss-Prot database or *Gallus gallus* proteome) had a smaller average length than those with known homologs (Fig. 6a & 6c). Similar results were found when this approach was used to assess genes predicted in the *R. marina* genome assembly (Edwards et al., 2018), and are indicative that these ‘unknown’ proteins are fragmented and lower quality predictions that may be due to underlying assembly issues with contiguity or frameshifting indels. The poorer quality could also reflect low stringency Maker2 gene predictions or homology based GeMoMa annotation of low-quality reference genes. The known proteins predicted by Maker2 (Fig. 6f) were of apparent lower quality than those reported by GeMoMa as indicated by their shorter lengths and lower protein ratios (Fig. 6e), which may be a result from a combination of incorrect gene predictions, and the high-quality reference homologs inflating quality scores of the GeMoMa annotation in comparison.

The known protein lengths were similar across the *S. vulgaris* vAU and vNA annotations (652 vs 650 aa), though there was a slightly larger difference in average unknown protein length (426 vs 407 aa). Although this increase in *S. vulgaris* vAU is very slight, it may indicate increased quality of unknown protein predictions in the vAU annotation, possibly due to the more Iso-Seq data mapping to the vAU genome (Fig. 4b) or the higher contiguity. Predicted genes were more commonly shorter than their closest reference protein hits, indicative there might still be some truncated gene predictions, consistent with the large number of assembly gaps. Nevertheless, the final annotation has a strong protein ratio peak around 1.0 for known proteins (Fig. 6b & 6d), indicating that the bulk of these predicted genes were of lengths similar to their Swiss-Prot homologs and hence deemed high quality.

Near identical assembly pipelines were used for the annotation of the two genome assemblies, with the resulting final gene count predictions comparable to other high-quality avian genomes and expected gene counts in eukaryote genomes. Both genome assembly versions reported similar final annotation statistics, with *S. vulgaris* vNA reporting slightly more predicted genes (Table 3), and a larger predicted gene coverage over the genome (59.09% gene coverage vs 55.23%), indicating this increase in predicted genes is not just a result of more overlapped predictions, though it could be a result of smaller assembly size and higher gene duplication (Fig. 5a).

## 5. Conclusion

This paper highlights the multifunctional use of species-specific transcript data, and the importance of diverse assessment tools in the assembly and assessment of reference genomes and annotations. We present a high-quality, annotated *S. vulgaris* vAU reference genome, scaffolded at the chromosome level. Alongside a second assembly, *S. vulgaris* vNA, these data provide vital resources for characterising the diverse and changing genomic landscape of this globally important avian. In addition to improving the completeness of gene annotation, we demonstrate the utility of long-read transcript data for genome quality assessment and assembly scaffolding. We also reveal some counter-intuitive behaviour of BUSCO genome completeness statistics, and present complementary two tools, BUSCOMP and SAAGA, which can identify and resolve potential artefacts, and inform assembly pipeline decisions.

## Author Contributions

Project conception: all authors

Sample Collection: KCS, SJW, MCB

Lab Work: KCS, YC, LAR, WCW

Data Analysis: KCS, RJE, YC, WCW

Program Development: RJE

Manuscript Writing: KCS, RJE

Manuscript Editing: All authors

## Acknowledgements

We thank non-author members of the Starling Genome Consortium for their support of this project including Wim Vanden Berghe. Thank you to Stella Loke, Annabel Whibley, and Mark Richardson for their guidance of Nanopore sequencing and analysis. Thank you to Daniel Selechnik for assistance with RNA extractions. Art credit (Fig. 7 illustration) to Megan Bishop. RJE was funded by the Australian Research Council (LP160100610 and LP18010072). DWB and YC acknowledge grant funding from the Human Sciences Frontier Programme (Grant RGP0030/2015). SM acknowledges Roslin Institute Strategic Grant funding from the UK Biotechnology and Biological Sciences Research Council (BB/P013759/1). LAR was supported by a Scientia Fellowship from UNSW.

## Data Accessibility and Programs

BUSCOMP documentation: https://slimsuite.github.io/buscomp/

Diploidocus documentation: https://slimsuite.github.io/diploidocus/

SAAGA documentation: https://slimsuite.github.io/saaga/

The data have been deposited with links to BioProject accession number PRJNA706841 in the NCBI BioProject database (https://www.ncbi.nlm.nih.gov/bioproject/). Bioproject raw data reviewer link is available at: https://dataview.ncbi.nlm.nih.gov/object/PRJNA706841?reviewer=op1k3u6792jbg8o4rddq138g9e

Genome accession JAGFZL000000000 is currently private on NCBI, but a reviewer copy is available here: https://drive.google.com/drive/folders/1MTExtlui-I-ziCIzwyBtgiP0NwHTg3IG?usp=sharing

Any scripts or metadata not covered by the above will be available on GitHub.

