## Supplementary Materials for "Transcript- and annotation-guided genome assembly of the European starling"

**Appendix 1: Genomic DNA sample collection, gDNA extraction, and sequencing**

A juvenile male starling (non-reproductive state) was collected (by a private landowner as part of local pest control measures) from Orange in New South Wales, Australia (33.2833° S, 149.1000° E), euthanized, immediately dissected, and the brain tissue was placed in RNALater, kept cold (3-4°C) for two weeks, and then stored for 2 months at -70°C. We extracted high molecular weight gDNA from the approximately 1g of brain tissue using the Gentra Puregene Tissue Kit (Qiagen, Hilden, Germany), as per the manufacturer's instructions. We checked the gDNA purity using a Nanodrop spectrophotometer (Thermo Fisher Scientific). We assessed fragment sizes using a Fragment Analyser (Millennium Science, Victoria, Australia) with an HS Large Fragment 50kb Kit (Agilent, California, USA). High molecular weight gDNA (1 ug) was prepared for 10x Chromium linked-read sequencing according to the manufacturer's recommended protocols. A 10x GEM library was barcoded using the Chromium Genome Reagent Kits (v2 Chemistry). The library was run on a single lane of a S4 flowcell and sequenced using the Illumina Hiseq X Ten sequencing platform (150 bp paired end reads).

High molecular weight gDNA (1 µg) was re-extracted for long read ONT (Oxford Nanopore Technologies, Oxford, United Kingdom) sequencing from the same brain tissue, using the 1D genomic DNA by ligation kit (SQK-LSK109, ONT) according to the standard protocol (with the inclusion of long fragment buffer). 500 ng of DNA was loaded onto two r9.4 minion flow cells and sequenced on an ONT MinION (Oxford Nanopore Technologies, Oxford, United Kingdom). Raw reads were converted to FASTA format, and filtered (guppy_basecaller parameters: min_qscore 7, filtlong parameters: min_mean_q 93, min_length 3000) with GUPPY (v.3.2.1) (Oxford Nanopore Technologies) using the high-accuracy flip-flop model (config file: dna_r9.4.1_450bps_hac.cfg).

### Appendix 2: Validation of Supernova genome size prediction using JELLYFISH

To confirm that the genome size estimate of 1.19 Gb produced by SUPERNOVA (v2.1.1) (Weisenfeld *et al.* 2017) was correct, we used k-mer frequency analysis through JELLYFISH (all k-mers counted) to manually confirm the estimated length of the genome based on the linked-read gDNA data. A k-mer histogram was produced using all the linked read gDNA raw data for an initial value of 20-mer based on approximated genome size of just above 1 Gb (Fig. S1). Counts for k-mer values of 7 or below (i.e., before the trough/red line) where removed as these are often attributed to either extremely rare reads, or random sequencing errors (Fig. S1). The genome size was estimated then by finding the total number of k-mers over all k values and dividing this by the point of mean coverage (Fig. S1, peak denoted by blue line; mean coverage = 32).

$$\text{Genome Length} = 35823112660 / 32$$
$$= 1{,}119{,}472{,}271 \text{ bp}$$

The final genome assembly size was used to find the using the genome size k-mer estimate.

$$1{,}049{,}838{,}585/1{,}119{,}472{,}271 = 93.78\% \text{ complete}$$

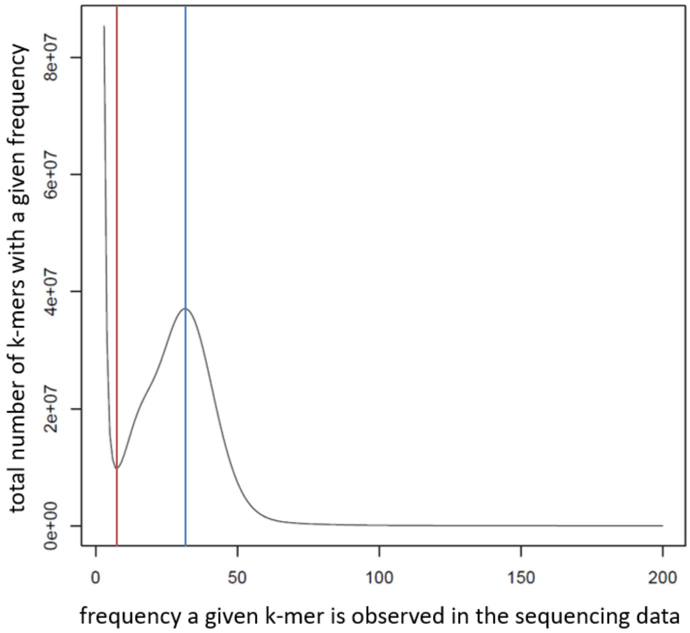


**Figure S1: Histogram of *Sturnus vulgaris* linked read gDNA Km-er counts** as calculated by JELLYFISH from k=7 to k=200. Vertical red line denotes histogram peak at k=37.

**Appendix 3: Transcriptome sample collection, RNA extraction, and sequencing**

An adult male European starling (non-breeding condition, black beak) was collected (by a private landowner as part of local pest control measures) via trapping from Wee Waa in New South Wales, Australia (30.2266° S, 149.4455° E) during late summer, with access to food and water. Within 24 hours of capture, the bird was euthanized, and we immediately dissected the brain, testes, and heart, and placed these in RNA-later, and kept cold (3-4$^{o}$C) for one week. We extracted RNA from whole brain, testes, and heart tissue using the RNeasy kit as per the manufacturer’s instructions (including DNA removal using RNase-free DNase treatment) (Qiagen, Hilden, Germany). We assessed RNA quality and purity using a Nanodrop spectrophotometer (Thermo Fisher Scientific, Victoria, Australia) and Qubit 4.0 (RNA IQ Assay Kit) (Invitrogen, California, United States). One cDNA synthesis reaction was carried out for each sample, with approximately 2 μg of total RNA used as starting material for the brain and heart and 1 μg for the testis. For the endpoint amplification, two parallel PCRs were carried out for each sample using the TeloPrime PCR Add-on Kit V2 (Lexogen) with 20 cycles and 2 μl of synthesised cDNA per reaction as template. The PCR products from the two reactions were pooled together and then split into two fractions, which were purified using 1x and 0.5x AMPure PB beads (Pacific Biosciences), respectively, and pooled at equal molarity. The heart and testis cDNA libraries were then combined at equal molarity. Three full-length cDNA libraries were constructed, one for each tissue, using the TeloPrime Full-Length cDNA Amplification Kit V2 (Lexogen) following the kit manual. For SMRTbell library prepartion, 740 ng and 677 ng of purified full-length cDNA were used for the brain and heart+testis libraries, respectively, as input for the SMRTbell Template Prep Kit 1.0 (Pacific Biosciences). Libraries were sequenced using full-length isoform transcript sequencing Iso-Seq program (PacBio, California, United States) on the PacBio Sequel 1 platform (software v6.0.0) using the V3 chemistry. The polymerase-bound libraries were sequenced on 1 SMRT Cell each with a 20 h movie time plus a 4 h pre-extension time using the Sequel Sequencing Kit 3.0 (PacBio, 101-597-900) and a Sequel SMRT Cell 1M v3 LR (PacBio, 101-531-001). Sequencing was performed at the Institute for Molecular Bioscience Sequencing Facility (University of Queensland; https://imb.uq.edu.au/sequencing-facility).

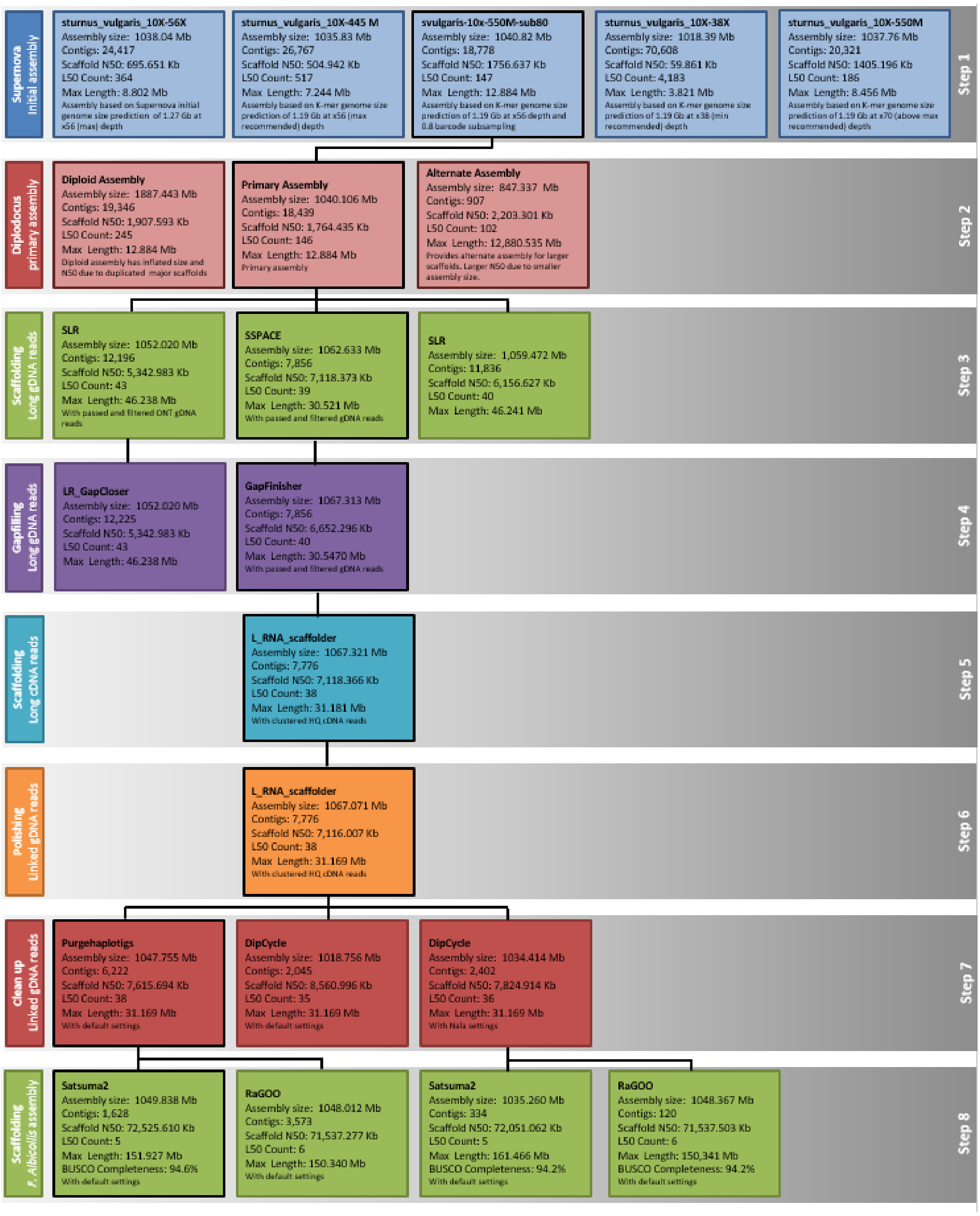


**Figure S2: Flow diagram of the *Sturnus vulgaris* vAU assembly process** covering initial assembly to final chromosome scaffolding, for a total of eight assembly steps; 1) primary assembly, 2) primary assembly, 3) ONT scaffolding, 4) gap-filling, 5) Iso-Seq scaffolding, 6) polishing, 7) cleanup, and 8) chromosome scaffolding.

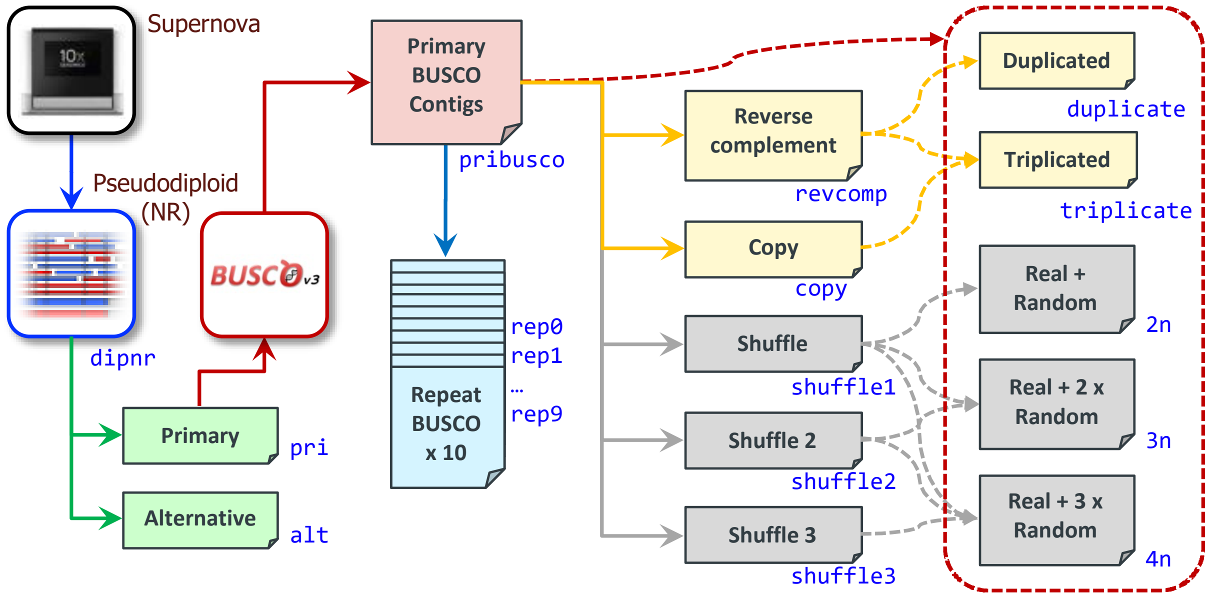


**Figure S3: BUSCO and BUSCOMP benchmarking datasets.** The SUPERNOVA pseudohap2 assembly was processed with DIPLOIDOCUS to generate a non-redundant pseudodiploid assembly ('dipnr') consisting of a primary ('pri') and alternative ('alt') assembly. BUSCO-containing scaffolds from the primary assembly were extracted into a reduced assembly ('pribusco') that was used to generate additional test data. **Duplication.** A reverse-complemented copy ('revcomp') was generated and combined with 'pribusco' to make a 100% duplicate assembly ('duplicate'). An additional direct copy ('copy') was then added to make a triplicated assembly ('triplicate'). **Assembly size.** Three randomly shuffled versions of 'pribusco' were generated ('shuffle1', 'shuffle2', 'shuffle3') and added in combination to `pribusco` to generate datasets of increasing assembly size without increasing duplication levels ('2n', '3n' and '4n'). **Reproducibility.** Ten straight repeats of the `pribusco` BUSCO run were also performed (`rep0` to `rep9`).

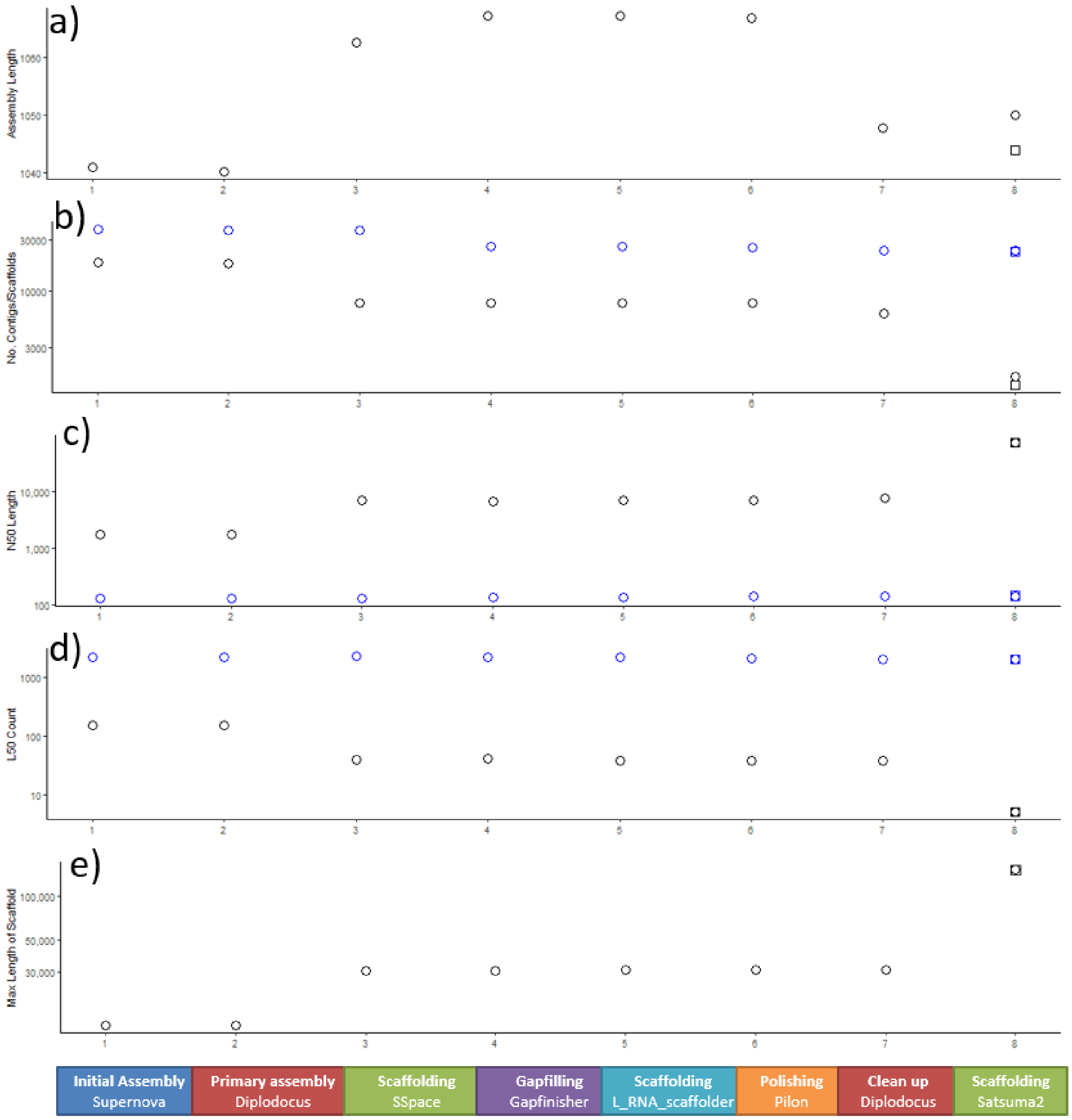


**Figure S4: *Sturnus vulgaris* vAU assessment statistics summary** for each of the eight assembly steps; 1) primary assembly, 2) primary assembly, 3) ONT scaffolding, 4) gap-filling, 5) Iso-Seq scaffolding, 6) polishing, 7) cleanup, and 8) chromosome scaffolding, with black indicating scaffolds and grey contigs, with vAU1.0 represented as circles, and vAU1.1 represented as squares.

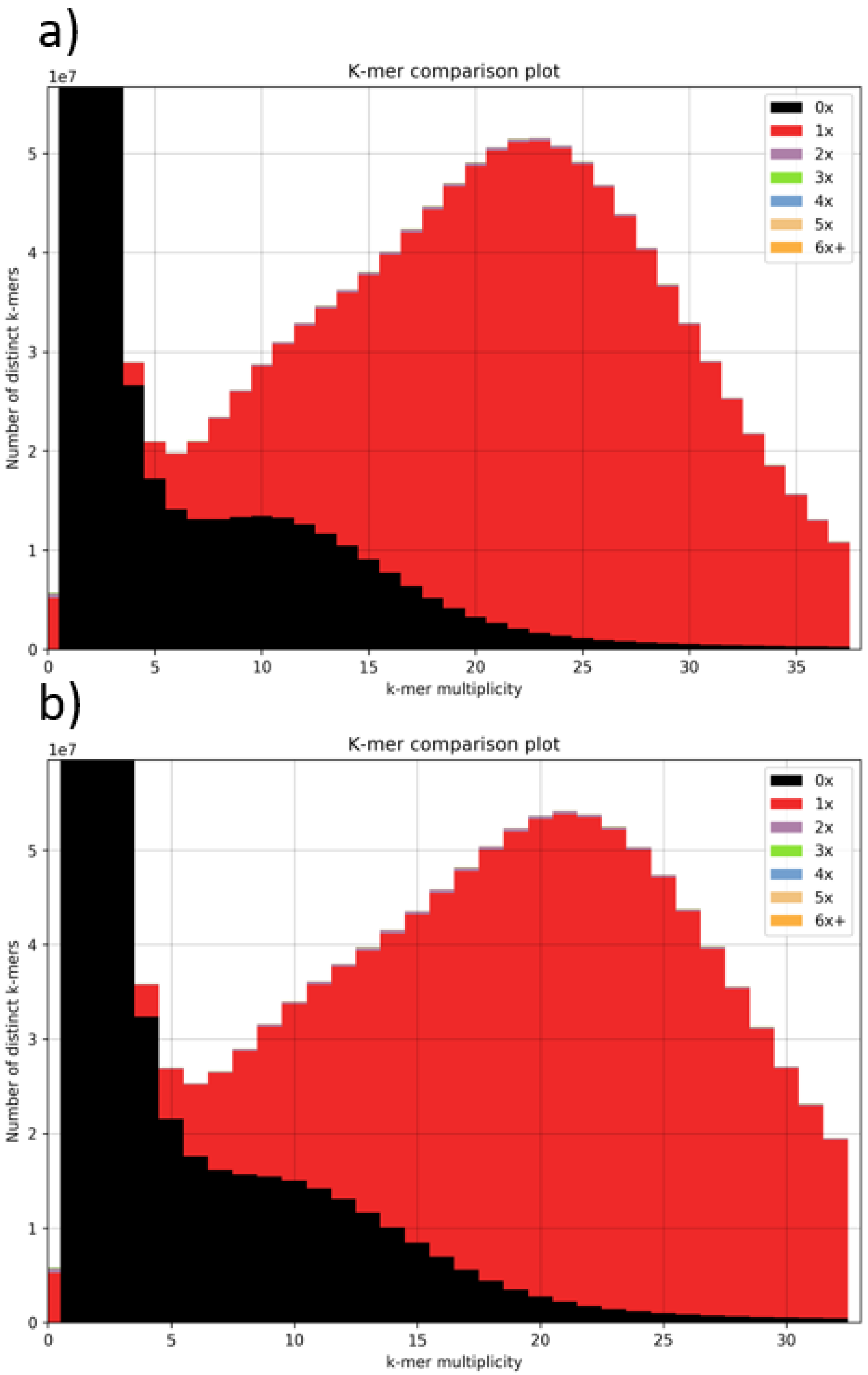


**Figure S5: KAT k-mer analysis of final *Sturnus vulgaris* AU assembly.** Plots depict read k-mer frequency distributions with different assembly copy numbers based on the 10X Chromium linked a) Reads 1 and b) Reads 2.

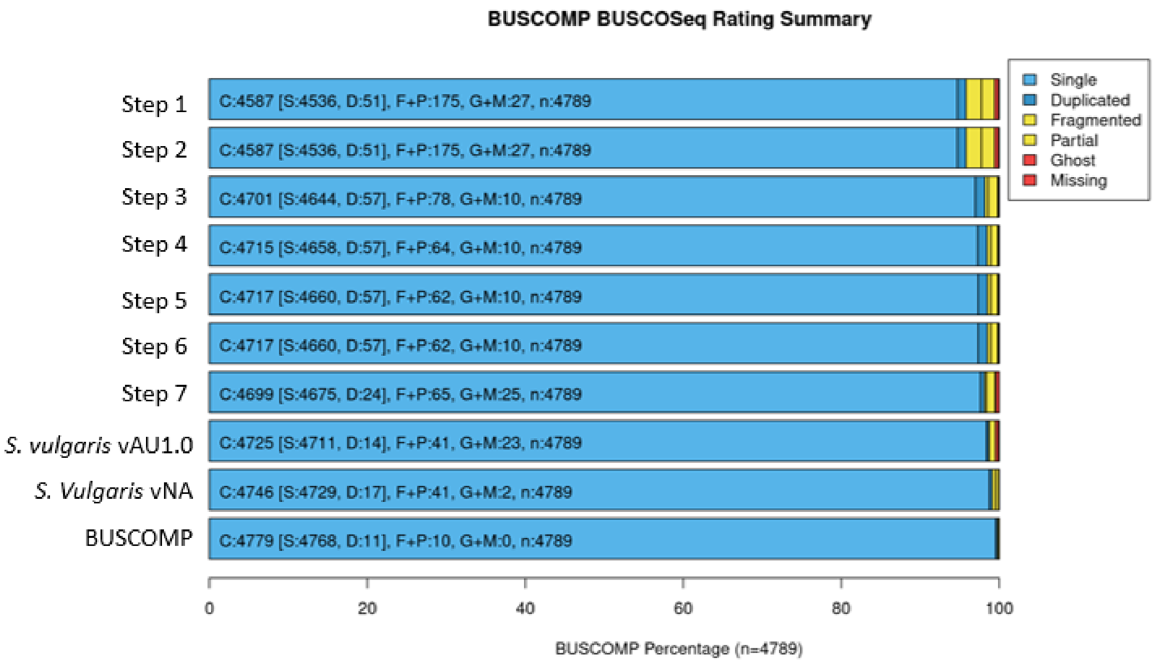


**Figure S6: BUSCOMP completeness** results for the 4,779 BUSCO genes identified as single copy and complete in one or more assembly stages, and including *S. vulgaris* vNA. The final BUSCOMP row compiles the best rating for each gene across all eight steps.

### Appendix 4: Assembly and annotation of the *Sturnus vulgaris* vNA genome version

**Materials and Methods:**

***Sample Collection, DNA extraction, and sequencing:*** The S. vulgaris DNA for shotgun sequencing was derived from an adult male (*Sturnus vulgaris*; bird ID 715) collected from North America. Total sequence genome input coverage on the Illumina HiSeq instrument was approx. 74x (39x fragments, and 35x 3kb) using a genome size estimate of 1Gb.

***Genome assembly and scaffolding**:* The combined sequence reads were assembled using ALLPATHS-LG software (Gnerre *et al.* 2011). This 1.0 version has been cleaned of contaminating contigs, and contigs 200 bp and less were removed.

***Genome Annotation and Functional Annotation**:* We annotated the *S. vulgaris* vNA genome using the same pipeline described in the main manuscript, with the only change being that a custom Augustus species profile was not trained for the Maker potion of the annotation.

**Results**

***Assembly:*** The assembly is made up of a total of 2361 scaffolds (including single contig scaffolds) with an N50 scaffold length of 3.6Mb (N50 contig length was 152kb). The assembly spans 1.01 Gb. The *S. vulgaris* reference genome can be downloaded on Genbank (Accession GCF_001447265.1).

***Annotation:*** The initial annotation produced by GEMOMA reported 20414 genes, with 77.2% BUSCO completeness (Figure S7). The initial MAKER2 annotation reported 15150 genes, and a BUSCO completeness of 97.6% (Figure S7). The merged final annotation reported a BUSCO completeness of 98.5% (Figure S7), containing 21,944 genes, and 81,714 mRNAs. There was an average of 11.8 exons and 10.8 introns per gene, with an average intron length of 3343. Of these, 1,122 were single exon genes and 2,018 were single-exon mRNAs. Predicted coding sequences made up 5.5% of the assembly, and 40.91% of the remaining sequences were unannotated.

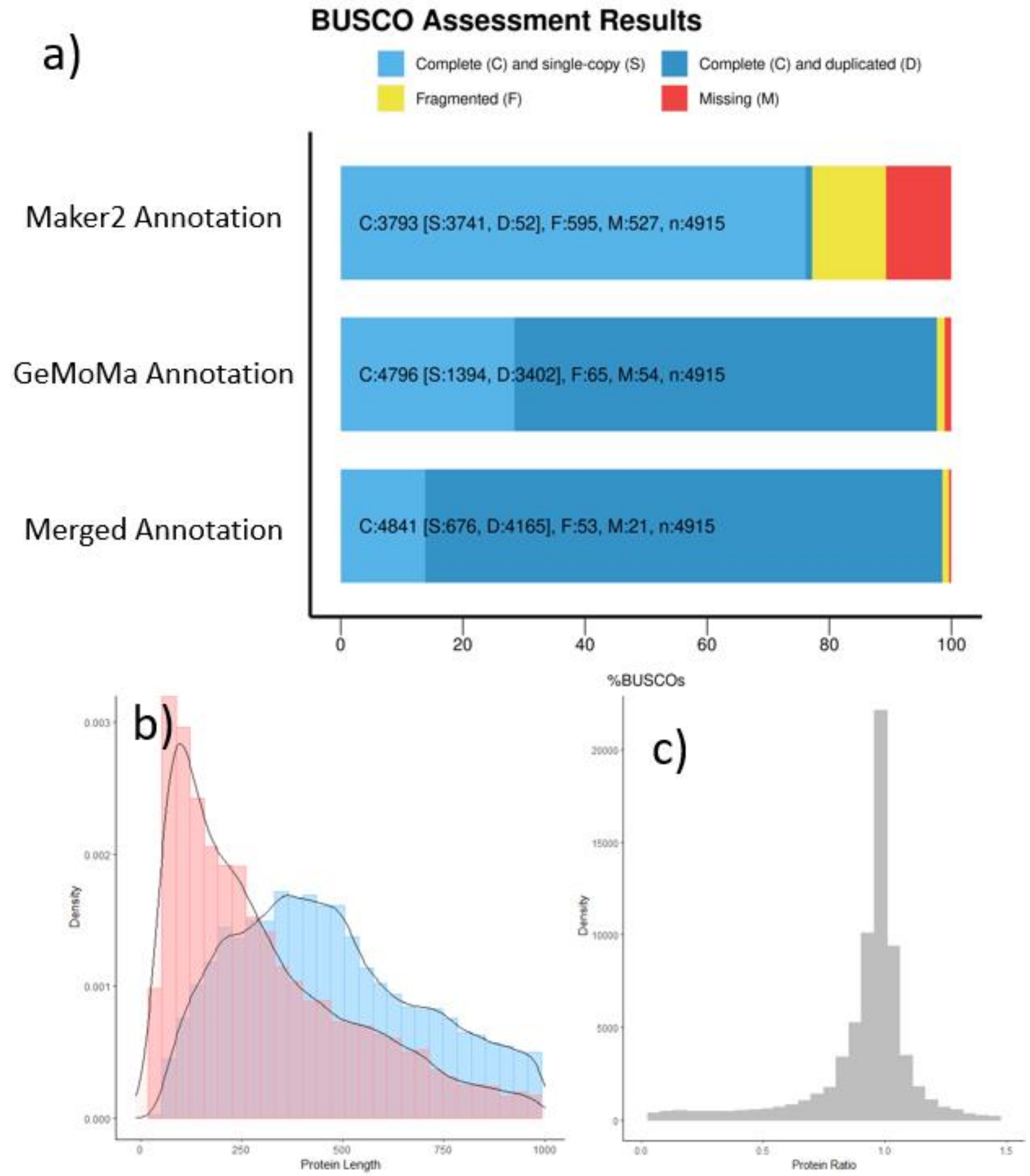


**Figure S7: Summary of *Sturnus vulgaris* vNA genome annotations**, with panel a) BUSCO assessments of initial Maker2 and GeMoMa assemblies, the final *S. vulgaris* vNU annotation (aves lineage). Panel b) depicts protein length of known and unknown proteins for the merged GeMoMa and Maker2 annotation. Panel d) depicts the protein ratio between output from SAAGA for all known proteins for the merged annotation (where a score close to 1 indicates a high-quality gene annotation, protein ratio calculated as annotated protein length / best Swiss-Prot reference protein length).

**Table S1: GeMoMa Ensembl reference species** used in GeMoMa annotation of *Sturnus vulgaris* vAU and vNA. Retrieved 18 Nov 2020.

| Common Name | Scientifiec Name | Ensembl Assembly |
|---|---|---|
| Eurasian sparrowhawk | *Accipiter nisus* | Accipiter_nisus_ver1.0 |
| Yellow-billed parrot | *Amazon collaria* | ASM394721v1 |
| Mallard | *Anas platyrhynchos* | ASM874695v1 |
| Pink-footed goose | *Anser brachyrhynchus* | ASM259213v1 |
| Swan goose | *Anser cygnoides* | GooseV1.0 |
| Greater spotted kiwi | *Apteryx haastii* | aptHaa1 |
| Little spotted kiwi | *Apteryx owenii* | aptOwe1 |
| Okarito brown kiwi | *Apteryx rowi* | aptRow1 |
| Golden eagle | *Aquila chrysaetos chrysaetos* | bAquChr1.2 |
| Burrowing owl | *Athene cunicularia* | athCun1 |
| Small tree finch | *Camarhynchus parvulus* | STF_HiC |
| Golden pheasant | *Chrysolophus pictus* | Chrysolophus_pictus_GenomeV1.0 |
| Blue tit | *Cyanistes caeruleus* | cyaCae2 |
| Emu | *Dromaius novaehollandiae* | droNov1 |
| Gouldian finch | *Erythrura gouldiae* | GouldianFinch |
| Flycatcher | *Ficedula albicollis* | FicAlb_1.4 |
| Chicken | *Gallus gallus* | GRCg6a |
| Median ground-finch | *Geospiza fortis* | GeoFor_1.0 |
| Bengalese finch | *Lonchura striata domestica* | LonStrDom1 |
| Turkey | *Meleagris gallopavo* | Turkey_2.01 |
| Great tit | *Parus major* | Parus_major1.1 |
| Ring-necked pheasant | *Phasianus colchicus* | ASM414374v1 |
| Common canary | *Serinus canaria* | SCA1 |
| African ostrich | *Struthio camelus australis* | ASM69896v1 |
| Zebra finch | *Taeniopygia guttata* | bTaeGut1_v1.p |
| White-throated sparrow | *Zonotrichia albicollis* | Zonotrichia_albicollis-1.0.1 |

**Table S2: NUMTfinder analysis of of final *Sturnus vulgaris* AU assembly,** identifying the location putative NUMT in the assembly.

| SeqName | Start | End | Strand | BitScore | Expect | Length | Identity | mtStart | mtEnd |
|---|---|---|---|---|---|---|---|---|---|
| SV_vAU_seq2 | 1.19E+08 | 1.19E+08 | - | 152 | 4.00E-33 | 302 | 216 | 6450 | 6751 |
| SV_vAU_seq4 | 47522543 | 47522606 | - | 72 | 1.00E-08 | 64 | 55 | 10558 | 10620 |
| SV_vAU_seq5 | 38040731 | 38040937 | - | 62 | 5.00E-06 | 210 | 142 | 15261 | 15470 |
| SV_vAU_seq5 | 1.01E+08 | 1.01E+08 | - | 198 | 1.00E-46 | 803 | 530 | 14706 | 15492 |
| SV_vAU_seq8 | 17525188 | 17525391 | + | 83 | 5.00E-12 | 208 | 145 | 15245 | 15451 |
| SV_vAU_seq8 | 25972974 | 25973368 | + | 189 | 5.00E-44 | 397 | 283 | 8702 | 9095 |
| SV_vAU_seq8 | 41187969 | 41188736 | + | 216 | 4.00E-52 | 814 | 545 | 14615 | 15413 |
| SV_vAU_seq9 | 17334457 | 17334507 | + | 70 | 3.00E-08 | 51 | 46 | 2525 | 2575 |
| SV_vAU_seq10 | 24554055 | 24554239 | - | 124 | 2.00E-24 | 192 | 144 | 12904 | 13094 |
| SV_vAU_seq11 | 355759 | 356192 | + | 291 | 6.00E-75 | 467 | 345 | 3622 | 4088 |
| SV_vAU_seq11 | 8642466 | 8642510 | + | 59 | 6.00E-05 | 45 | 40 | 2088 | 2132 |
| SV_vAU_seq11 | 30396630 | 30396961 | - | 91 | 1.00E-14 | 338 | 227 | 14615 | 14944 |
| SV_vAU_seq12 | 2721351 | 2722141 | - | 537 | 8.00E-149 | 838 | 617 | 2753 | 3578 |
| SV_vAU_seq12 | 4344254 | 4344336 | - | 65 | 1.00E-06 | 83 | 64 | 14345 | 14427 |
| SV_vAU_seq14 | 1657092 | 1657312 | - | 192 | 4.00E-45 | 221 | 175 | 8819 | 9039 |
| SV_vAU_seq14 | 4520796 | 4521008 | - | 133 | 3.00E-27 | 220 | 161 | 12804 | 13023 |
| SV_vAU_seq16 | 788033 | 788121 | + | 62 | 5.00E-06 | 89 | 67 | 10565 | 10653 |
| SV_vAU_seq24 | 2478314 | 2478498 | - | 69 | 3.00E-08 | 186 | 130 | 16408 | 16590 |
| SV_vAU_seq26 | 2392491 | 2392570 | - | 86 | 4.00E-13 | 80 | 67 | 14106 | 14185 |
| SV_vAU_seq26 | 2589746 | 2590569 | + | 159 | 2.00E-35 | 894 | 580 | 13616 | 14501 |
| SV_vAU_seq26 | 2696879 | 2697117 | + | 125 | 5.00E-25 | 240 | 173 | 16409 | 16646 |
| SV_vAU_seq31 | 37052129 | 37052981 | - | 829 | 0 | 863 | 705 | 11644 | 12504 |
| SV_vAU_seq31 | 42850393 | 42851068 | + | 369 | 3.00E-98 | 680 | 498 | 15153 | 15823 |
| SV_vAU_seq31 | 43349606 | 43349796 | - | 68 | 1.00E-07 | 193 | 133 | 4484 | 4675 |
| SV_vAU_seq31 | 62603293 | 62603703 | - | 84 | 2.00E-12 | 451 | 288 | 10162 | 10607 |
| SV_vAU_seq31 | 62603804 | 62604202 | + | 123 | 2.00E-24 | 425 | 282 | 4203 | 4622 |
| SV_vAU_seq31 | 68129474 | 68129742 | - | 96 | 2.00E-16 | 275 | 187 | 8025 | 8289 |

**Table S3: Assembly statistics summary** for sequential assembly steps of *Sturnus vulgaris vAU1.0*

| | Step 1: Assembly | Step 2: Primary Assembly | Step 3: Long gDNA read Scaffolding | Step 4: Gapfilling | Step 5: Long cDNA read Scaffolding | Step 6: Polishing | Step 7: Tidy | Step 8: Chromosome Scaffolding |
|---|---|---|---|---|---|---|---|---|
| **Assembly continuity statistics** | | | | | | | | |
| Total number of sequences | 18,778 | 18,439 | 7,856 | 7,856 | 7,776 | 7,776 | 6,222 | 1,628 |
| Total length of sequences | 1,040,824,271 | 1,040,106,492 | 1,062,633,441 | 1,067,313,776 | 1,067,321,776 | 1,067,071,200 | 1,047,755,039 | 1,049,838,585 |
| Min. length of sequences | 1,000 | 1,000 | 1,000 | 977 | 977 | 842 | 917 | 927 |
| Max. length of sequences | 12,884,419 | 12,884,419 | 30,521,271 | 30,547,435 | 31,181,295 | 31,169,695 | 31,169,695 | 151,927,750 |
| Mean length of sequences | 55,427.86 | 56,407.97 | 135,263.93 | 135,859.70 | 137,258.46 | 137,226.23 | 168,395.22 | 644,864.00 |
| Median length of sequences | 2,121 | 2,153 | 2,433 | 2,433 | 2,395 | 2,394 | 2,199 | 1,337 |
| N50 length of sequences | 1,756,637 | 1,764,435 | 7,118,373 | 6,652,296 | 7,118,366 | 7,116,007 | 7,615,694 | 72,525,610 |
| L50 count of sequences | 147 | 146 | 39 | 40 | 38 | 38 | 37 | 5 |
| Total number of contigs | 37,718 | 37,354 | 36,809 | 26,531 | 26,531 | 25,946 | 23,815 | 23,815 |
| Contig N50 length of sequences | 132,941 | 132,973 | 131,172 | 138,163 | 138,163 | 143,478 | 146,413 | 145,864 |
| Contig L50 count of sequences | 2,176 | 2,174 | 2,239 | 2,151 | 2,151 | 2,079 | 2,023 | 2,030 |
| GC content | 41.64% | 41.64% | 41.68% | 41.81% | 41.81% | 41.82% | 41.73% | 41.73% |
| N bases | 6,838,110 (0.66%) | 6,835,790 (0.66%) | 23,897,712 (2.25%) | 11,590,876 (1.09%) | 11,598,876 (1.09%) | 11,396,143 (1.07%) | 11,158,567 (1.06%) | 13,242,113 (1.26%) |
| Gap (10+ N) length | 6,838,110 (0.66%) | 6,835,790 (0.66%) | 23,895,932 (2.25%) | 11,590,274 (1.09%) | 11,598,274 (1.09%) | 11,395,574 (1.07%) | 11,158,028 (1.06%) | 13,241,574 (1.26%) |
| Gap (10+ N) count | 18,940 | 18,915 | 28,953 | 18,675 | 18,755 | 18,170 | 17,593 | 22,187 |
| **BUSCO statistics** | | | | | | | | |
| Complete and single copy | 4,470 | 4,470 | 4,536 | 4,541 | 4,537 | 4,644 | 4,579 | 4,595 |
| Complete and duplicate | 95 | 95 | 101 | 99 | 99 | 98 | 61 | 54 |
| Fragmented Busco | 219 | 219 | 173 | 169 | 170 | 168 | 164 | 154 |
| Missing Busco | 131 | 131 | 104 | 143 | 107 | 103 | 110 | 112 |
| **Iso-Seq mapping statistics** | | | | | | | | |
| ISO-SEQ:Non-mapped transcripts | 264 | 264 | 267 | 244 | 247 | 246 | 246 | 241 |

| | | | | | | | | |
|---|---|---|---|---|---|---|---|---|
| ISO-SEQ: Mapped transcripts (Quality = 60) | 32,685 | 32,693 | 32,600 | 32,565 | 32,572 | 32,578 | 32,898 | 32,864 |
| **BUSCOMP Statistics** | | | | | | | | |
| BUSCOMP: Complete and single copy | 4,530 | 4,530 | 4,632 | 4,640 | 4,642 | 4,642 | 4,660 | 4,702 |
| BUSCOMP: Complete and duplicate | 48 | 48 | 55 | 55 | 55 | 55 | 22 | 13 |
| BUSCOMP: Fragmented and Partial | 129 | 129 | 38 | 30 | 28 | 28 | 29 | 12 |
| BUSCOMP: Ghost and Missing | 20 | 20 | 2 | 2 | 2 | 2 | 16 | 0 |
| **SAAGA statistics** | | | | | | | | |
| SAAGA: mean protratio | 0.919338 | 0.919562 | 0.936762 | 0.937015 | 0.938079 | 0.937305 | 0.940878 | 0.946452 |
| SAAGA: protratio_median | 0.998066 | 0.998069 | 0.998674 | 0.998720 | 0.998779 | 0.998773 | 0.998924 | 0.999204 |
| SAAGA: protratio_sd | 0.216565 | 0.216359 | 0.192846 | 0.193475 | 0.192076 | 0.192694 | 0.186809 | 0.178068 |
| SAAGA: duplicity | 0.918160 | 0.917762 | 0.913638 | 0.914098 | 0.913664 | 0.915717 | 0.897704 | 0.897401 |
| SAAGA: mean F1 score | 0.906657 | 0.906810 | 0.922552 | 0.922834 | 0.923881 | 0.923425 | 0.927146 | 0.931119 |
| SAAGA: mean_f1 | 0.598339 | 0.598838 | 0.595627 | 0.586467 | 0.587486 | 0.588122 | 0.593248 | 0.598555 |
| **KAT statistics** | | | | | | | | |
| KAT: R1 reads | 96.54 | 96.54 | 96.54 | 96.61 | 96.61 | 96.73 | 96.24 | 96.69 |
| KAT: R2 reads | 95.01 | 95.77 | 95.77 | 95.84 | 95.84 | 95.96 | 95.46 | 95.92 |

**Table S4: Read number and length after Iso-Seq analysis** from full-length transcriptome sequencing from *Sturnus vulgaris* brain, and heart and testes pooled.

| | **Brain** | **Heart + Testis** |
|---|---|---|
| **Raw Data** | | |
| Polymerase Read Bases | 39,550,574,401 | 30,329,883,755 |
| Polymerase Reads | 648,290 | 600,764 |
| Polymerase Read Length (Mean) | 61,008 | 50,486 |
| Polymerase Read N50 | 125,257 | 101,226 |
| Subread Length (mean) | 1,880 | 1,556 |
| Subread N50 | 2,069 | 1,719 |
| Insert Length (mean) | 2,985 | 2,625 |
| Insert N50 | 3,225 | 2,866 |
| **Primer removal + demultiplexing** | | |
| ZMWs input | 483167 | 446827 |
| ZMWs above all thresholds | 451260 (93%) | 419191 (94%) |
| ZMWs below any threshold | 31907 (7%) | 27636 (6%) |
| **Refine** | | |
| Number of reads | 446838 | 414411 |
| Number of reads (polya) | 445670 | 413023 |
| **Clustered** | | |
| High quality | 33454 | |
| High quality: mean length | 2005.79 | |
| Low quality | 157 | |
| **TAMA Collapse** | | |
| Non redundant transcripts | 28448 | |
| Non redundant transcripts: mean (bp) | 2014.43 | |
